# The influence of liver fluke infection on production in sheep and cattle: a meta-analysis

**DOI:** 10.1101/2020.07.29.227074

**Authors:** Adam D. Hayward, Philip J. Skuce, Tom N. McNeilly

**Affiliations:** Moredun Research Institute, Pentlands Science Park, Bush Loan, Penicuik, Midlothian, EH26 0PZ

**Keywords:** *Fasciola hepatica*, *Fasciola gigantica*, trematode, production, disease, systematic review

## Abstract

Liver flukes (*Fasciola* spp) are important parasites of sheep and cattle across the world, causing significant damage to animal health and productivity due to both acute and chronic infection. Many comprehensive reviews have discussed the results of decades of research into the impact of fluke infection on livestock performance traits such as weight gain and milk production. While fluke are considered to be important, there have been no attempts to collate previous research in a quantitative manner, and nor has there been an attempt to determine why some studies find substantial effects of fluke while others conclude that effects of fluke on animal performance are negligible. In this study, we used meta-analysis to provide quantitative estimates of the impact of liver fluke on animal performance, and to identify elements of study design that influence the conclusions of such studies. A literature search provided 233 comparisons of performance in “fluke-infected” and “uninfected” animals. We standardized these data as log response ratios and calculated effect size variances in order to weight studies by their sample size and accuracy of their estimates. We performed multi-level meta-analysis to estimate effects of fluke infection in five traits: daily weight gain (N = 77); live weight (N = 47); carcass weight (N = 84); total weight gain (N = 18) and milk production (N = 6). There were significant negative effects of fluke infection on daily weight gain, live weight and carcass weight (9%, 6% and 0.6% reductions in performance, respectively), but not total weight gain or milk production. We then used mixed-effects meta-analysis to estimate the impact of moderator variables, including host, fluke, and study design factors, on study outcomes. We found that, in general, studies that gave experimental infections found generally larger effects of fluke than observational or drug studies; younger animals were more likely to suffer the effects of fluke infection on daily weight gain; and that effects on live weight increased across the course of an experiment. Our results provide the first quantitative estimate of the importance of liver fluke on performance across studies and highlight the elements of study design that can influence conclusions. Furthermore, our literature search revealed areas of research into liver fluke that could be the subject of greater effort, and types of study that could form the basis of future meta-analyses.

## INTRODUCTION

Liver flukes (*Fasciola* spp) are amongst the most important helminth parasites of domestic sheep and cattle worldwide, causing significant financial losses to producers (Schweizer et al., 2005). They have a typical trematode parasite life-cycle: adults inhabit the host liver and bile duct system and produce eggs, which are shed in faeces. Miracidia develop within the eggs and then hatch and search for a mud snail (typically, *Galba trunculata*) intermediate host, which they penetrate, undergoing multiplication before emerging as cercariae, which encyst on vegetation as the infectious metacercariae (cysts). These are ingested by the host while grazing; immature fluke then emerge and migrate through the intestinal wall to the liver, where they develop into adults around 10-12 weeks after ingestion of cysts (Skuce and Zadoks, 2013). Liver fluke can cause acute disease associated with migration of immature fluke, which can lead to death in severe cases, especially in sheep, and chronic disease caused by the blood-feeding activity of the adults, which can live as long as the host (Kaplan, 2001). Control of liver fluke remains a challenge in all areas of the world: vaccine development has been difficult, due in part to the lack of a robust protective host immune response and a lack of understanding of which antigens to target (Molina-Hernández et al., 2015; Toet et al., 2014); increasing flukicide resistance (Brockwell et al., 2013; Kamaludeen et al., 2019; Novobilský and Höglund, 2015); the clonal amplification of the parasite in the intermediate snail host (Beesley et al., 2018); and the ability of wildlife reservoir hosts to disseminate the parasite (French et al., 2019).

A large number of influential reviews have collated decades of research into the effects of fluke infection on animal performance (Charlier et al., 2013; Dargie, 1980, 1987; Elelu and Eisler, 2018; Skuce and Zadoks, 2013). Empirical work has demonstrated statistically significant and sometimes substantial effects of liver fluke infection on traits including carcass weight (Sanchez-Vazquez and Lewis, 2013), carcass conformation or fatness (Bellet et al., 2016; Sanchez-Vazquez and Lewis, 2013), age at slaughter (Mazeri et al., 2017), weight gain (Chick et al., 1980; Genicot et al., 1991; Hope Cawdery et al., 1977; Loyacano et al., 2002; Sykes et al., 1980), milk production (El-Tahawy et al., 2017; May et al., 2020), as well as the financial costs associated with condemnation of infected livers (Nyirenda et al., 2019). Other studies, however, have found no support for effects of liver fluke infection on performance traits including carcass weight (Bellet et al., 2016; Charlier et al., 2009; Molina et al., 2005), weight gain (Bossaert et al., 2000; Echevarria et al., 1992; Forbes et al., 2015) and milk production (May et al., 2019; Randall and Bradley, 1980). These studies vary in the direction and magnitude of effects of fluke, but also in their characteristics. Firstly, there is biological variation: in parasite species (*F. hepatica* typically in temperate regions and *F. gigantica* in the tropics) and in animal breed, age and sex. There is also variation in study design: some studies compare naturally infected with uninfected animals, some compare control with experimentally-infected animals, and others compare control with flukicide-treated animals. Finally, effects may vary with location due to climatic conditions, and with the time since initial infection, as it takes around 8-12 weeks for fluke to migrate to the liver and mature (Kaplan, 2001). As such, we lack knowledge on (1) the overall impact of fluke across studies and (2) the effects of study characteristics on outcome.

In this study, we present a meta-analysis of the impact of liver fluke infection on performance in sheep and cattle. Meta-analysis aims to collate data from published and unpublished sources addressing the same question and then puts data from these studies onto a standardized scale (“effect sizes”), enabling statistical analysis of the overall effect and causes of variation in outcomes (Gurevitch et al., 2018). The term “meta-analysis” was coined in 1976 (Glass, 1976) and the techniques were quickly embraced by medical and social sciences, with studies in ecology and evolution beginning in the early 1990s (Lau et al., 2013). Only more recently has this approach been applied in veterinary science (Lean et al., 2009). We followed the PICO (population; intervention; comparator; outcome) approach to formulate our research questions (Stewart et al., 2013), aiming to compare different measures of weight and milk production in sheep and cattle infected with fluke against those designated as uninfected. We first estimated an overall effect size using random-effects meta-analysis and then assessed the impact of biological and study design factors (“moderators”) on study outcomes. Our results reveal that fluke infection has a particularly strong influence on weight gain, and that animal age and experimental design are important factors influencing study outcome.

## METHODS

### Literature search

We searched the scientific literature in order to identify studies that investigated the impact of liver fluke on performance of sheep and cattle. A *Web of Science* search was conducted on 18/10/2019 with the search terms (Fasciola OR fluke) AND (Cattle OR cow* OR calf OR calves OR sheep) AND (producti* OR weight OR grow* OR milk OR performance OR fertility OR carcas*) and the search yielded 662 papers. To these, we added all papers cited in a number of influential reviews on the impacts of fluke on livestock productivity (Charlier et al 2014A; Charlier et al 2014B; Dargie 1987; Elelu & Eiser 2018; Skuce & Zadoks 2013; Dargie 1980). We then added all papers citing these articles using the *Publish or Perish* software (Harzing 2016). Finally, we added a paper by da Costa et al (2019) that was published in late October 2019. We also added data provided by Scotbeef Ltd (Hayward et al., in prep) and McIntosh-Donald Ltd (Skuce et al., in prep). This resulted in a total of 1582 data sources.

We reviewed the titles and abstracts of these publications, sifting out publications that were clearly unsuitable for a variety of reasons (**Figure S1**). Once duplicates were removed, this initial sift resulted in 106 publications that were fully reviewed. Specifically, we searched for papers that compared performance in groups of animals that were infected with fluke (naturally or through experimental infection) versus animals that were uninfected (naturally or through flukicide treatment). We collected data on the mean, standard deviation, standard error and number of animals with performance measured in each group. Where data were presented in figures but not in tables or text, we used the R package ‘metaDigitise’ (Pick et al 2018) to extract data. Where it seemed that relevant data may been collected but not reported in the publication or supporting information, we contacted authors in order to request data. Once the full review was complete and unsuitable publications removed (**Figure S2**), our final dataset consisted of 233 effect sizes from 28 sources (**Table 1**).

**Table 1.**
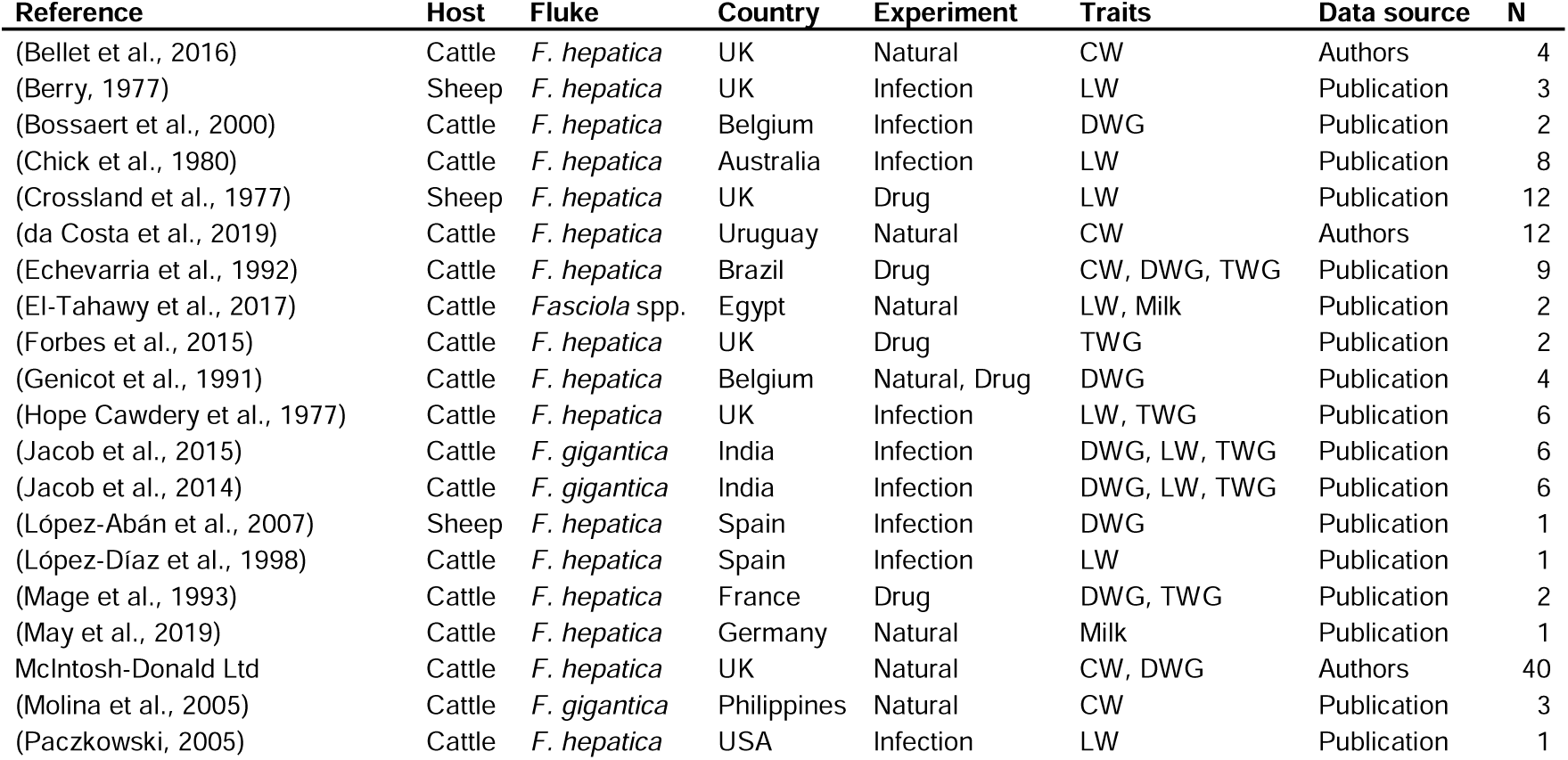
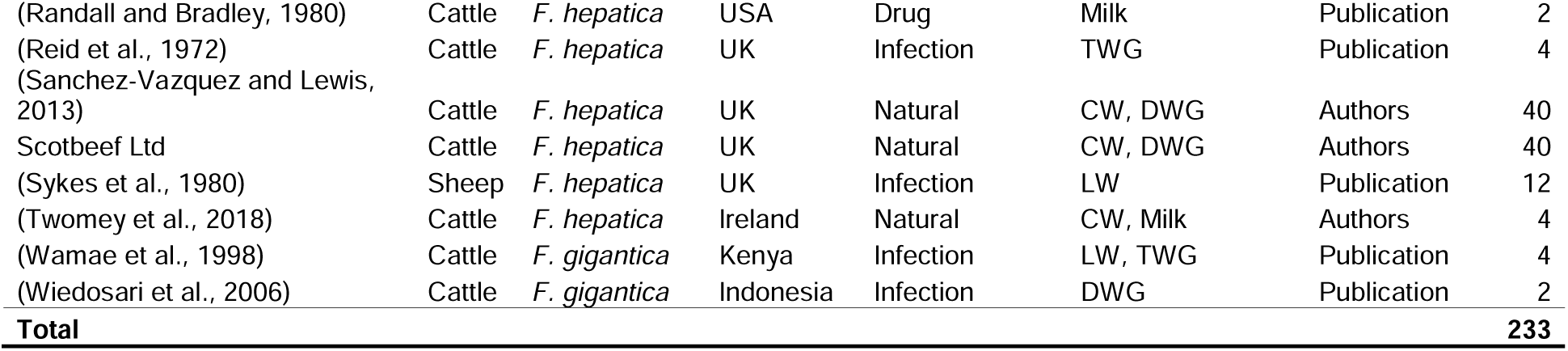
A summary of the studies from which data was used for the meta-analysis. ‘Experiments’ are as described in the main text, while traits are as follows: CW = carcass weight; DWG = daily weight gain; LW = live weight; milk = milk production; TWG = total weight gain. The data source was either the publication (taken from the text, tables of figures in the publication) or the authors (send upon request by authors). ‘N’ is the number of effect sizes contributed by each study.

### Data synthesis

We analysed the influence of liver fluke infection on five performance traits, as follows:

#### Daily weight gain

the calculated average increase in body weight per day. In some studies – mostly experimental – this is live weight gain (weight of the live animal divided by time in days), but in abattoir studies, this is usually dead weight gain (carcass weight divided by age in days). These effects are considered the same trait due to (1) the generally close correlation between live weight and carcass weight and (2) the way in which we tested for effects of study design in our analysis.

#### Live weight

the weight of the live animal, generally reported multiple times across the course of experimental studies.

#### Carcass weight

the weight of the animal’s carcass in abattoir studies.

#### Total weight gain

the amount of weight gained by the animal from the start of an experiment to the time of measurement.

#### Milk production

the weight of milk produced, generally expressed as a daily rate.

Our included studies reported raw mean performance in animals that were deemed to be fluke-infected versus animals that were deemed to be uninfected, plus sample sizes for both groups and standard deviation. Where standard errors were reported, we estimated the standard deviation as 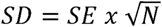. We calculated the ratio of means between fluke-infected and uninfected animals as our standardized measure of effect size and used log-transformation to normalize the values. Thus, we calculated log response ratio as 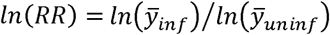, where 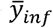 and 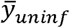 are the means of the trait *y* in infected and uninfected animals respectively. We calculated the variance for each effect as 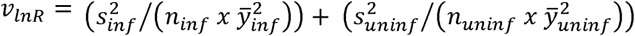 where *s*^2^, *n*, and 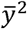 are the squared standard deviation, sample size and squared mean for infected or uninfected individuals (Koricheva et al 2013).

The 233 effect sizes were unevenly distributed across our 28 sources, with 4 studies contributing just one effect size, 5 contributing more than 10 and 3 studies contributing 40 effect sizes each. These last three were abattoir studies where we were provided with raw data by the authors or the abattoirs themselves and we calculated breed- and sex-specific statistics for the ten commonest breeds in each data set, in order to better account for these factors and to maximise our number of effect sizes. Multiple effect sizes were contributed for a number of reasons including comparisons being made across many weeks of an experimental study; fluke status being manipulated as well as another experimental treatment such as diet; data being subdivided by sex and/or breed. We categorized our effect sizes according to a number of biological and experimental factors, many of which we included in our meta-regression analyses (see below).

#### Host species

our data contained more effect sizes from cattle (205, 88%) than sheep (28, 12%).

#### Host breed

data came from 4 breeds of sheep and 29 breeds of cattle. Among these were crosses and cattle denoted simply as “dairy”, “beef” or “mixed”, each of which was included as a separate breed in our analyses.

#### Host sex

our data contained 88 effect sizes from females (38%), 106 from males (45%), 30 from mixed-sex groups (13%) and 9 where animal sex was not recorded (4%).

#### Host age group

animals were divided into three age categories, namely adults (12% of effect sizes), young (35%) and mixed (53%). For cattle, young animals were ≤ 12 months of age and adults ≥ 23 months of age. For sheep, young animals were ≤ 12-13 months of age and adults were year old (yearlings) or older.

#### Parasite

the data were dominated by *F. hepatica*, which accounted for 90% of effect sizes, with 9% contributed by *F. gigantica* and 1% from a single study recording a mixed *Fasciola* species burden.

#### Experimental design

29 effect sizes (12%) came from 6 studies where flukicidal drugs were used to remove fluke from an experimental group, performance in which was compared with a control group. In the drug-treated (i.e. “uninfected”) groups, the maximum mean fluke FEC was 3 eggs/gram and the maximum live fluke burden at post-mortem was 3. While some of these animals defined as “uninfected” clearly carried fluke, we consider these burdens low enough as to be negligible. These studies were denoted as “drug”. 56 (24%) effect sizes were from studies (denoted as “infection”) where animals were experimentally infected with fluke and compared with uninfected controls. The remaining 148 (64%), denoted “natural”, were largely from abattoirs and animals that acquired infection (or not) naturally.

#### Week post-infection

for experimental studies, data were collected from 4-54 weeks post-infection.

#### Absolute latitude

our effect sizes were predominantly from studies conducted in the UK (64%) with others coming from studies conducted in the Americas (12%), Asia (9%), Europe (8%), Australasia (4%), and Africa (3%). Absolute latitude ranged from 1.25-57.3.

### Multi-level meta-analysis

Meta-analyses were performed using the rma.mv function of the R package ‘metafor’ (Viechtbauer 2010), version 2.1-0. We performed separate random-effects meta-analyses in order to estimate the mean effect of fluke infection on five traits: daily weight gain, live weight, carcass weight, total weight gain and milk yield.

#### Global effects

First, we determined the mean effect of fluke infection on each of the traits. We used multi-level analyses: in order to account for non-independence of effect sizes derived from the same studies and from animals of the same breed, we fitted random effects of study and breed, as well as an observation-level random effect in order to estimate the residual variance.

#### Meta-regression

Next, we used a meta-regression approach in order to investigate whether the effect size depended upon a number of factors related to host and parasite biology or study design. These included host species (cattle or sheep), host sex, host age group (adult, young, or mixed), parasite species (*F. hepatica* or *F. gigantica*), study design (observational study of natural infections, infection experiment, drug treatment experiment), absolute latitude (continuous), and week post-infection. Each of these moderating variables was investigated in a separate meta-regression model. In each model, we determined whether the moderator was supported (i.e. whether the effect size varied according to the moderator) using Wald-type chi-square tests (Q_M_). We examined whether each level within categorical moderators was significantly different from zero using z-tests. We did not test all of the moderators for each of the five traits, since not all traits had variation in all moderators (e.g. daily weight gain data all came from cattle and so it was not possible to test the ‘host species’ moderator).

Details on the terms fitted for each trait are shown in **Table 2**.

**Table 2.** Summary of parameters included in meta-analysis of six production traits. ‘N’ and ‘studies’ refers to the maximum number of effect sizes available for each trait, and the number of studies measuring each trait.

| **Trait** | **N** | **Studies** | **Random effects** | **Moderators tested in meta-regression** |
| --- | --- | --- | --- | --- |
| Daily weight gain (DWG) | 77 | 11 | Study + Residual | Parasite, Age, Experiment, Sex, Latitude, Week |
| Live weight | 47 | 11 | Study + Residual | Parasite, Host, Age, Experiment, Sex, Latitude, Week |
| Carcass weight | 84 | 8 | Study + Residual | Parasite, Age, Experiment, Sex, Latitude |
| Total weight gain (TWG) | 18 | 8 | Study + Breed + Residual | Parasite, Experiment, Sex, Latitude, Week |
| Milk production | 6 | 4 | Residual | Experiment, Latitude |

### Analysis of heterogeneity and bias

Heterogeneity in effect sizes may be generated through variation between studies and variation within studies. In order to quantify this, we calculated the proportion of total variance that was due to variation in effect sizes (the *I*^*2*^ statistic), where the remainder is accounted for by sampling error. Since we fitted random effects to account for expected similarity between effect sizes from the same study and from the same breed of animal, we calculated modified *I*^*2*^ (Nakagawa & Santos 2012). Hence, for each meta-analysis, we calculated *I*^*2*^ values for the between-study effect and between-breed effect (where these were fitted (**Table 2**), and the residual effect.

The causes of bias in meta-analyses can include publication bias (the possibility that studies finding non-significant effects will not be published) and changes in effect size across time, e.g. if literature searches are less able to sample early sources. To test for publication bias, we generated funnel plots and performed regression of meta-analytic model residuals of each effect size (corrected for random effects and significant moderators) on the variance in each effect size (Egger, 1997; Nakagawa & Santos 2012). Where the intercept was significantly different from zero, we concluded that there was significant bias (Nakagawa & Santos, 2012). Where our data included both published and unpublished studies, we performed this test on both the full dataset, and then on the published studies only.

Finally, for each meta-analysis, we tested the effect of year of study as a continuous moderator in order to determine whether there has been a linear change in effect size across time, assessing significance using Wald-type chi-square tests (Q_M_).

## RESULTS

### Daily weight gain

Multi-level meta-analysis of daily weight gain revealed an overall negative effect of fluke infection (*β*_*global*_ = −0.0981, 95%CI = −0.1554 – −0.0408, z = −3.36, P < 0.001), suggesting that infected animals gained 9.5% less weight per day than uninfected animals (Figure 1). Meta-regression analyses revealed that the moderators of parasite species, experimental type, sex, latitude and week had no influence on the effect size (Table 3), but that there was a significant effect of the moderator of age (Q_M_ = 5.29, P = 0.021), suggesting that young animals, but not mixed-aged groups, experienced negative effects of fluke. Our results also revealed that there were negative effects of fluke both in animals infected with *F. gigantica* and *F. hepatica*, studies that used experimental infections but not drug studies or studies using natural infections, and in both sexes (Figure 1). Total variation in effect sizes was high (*I*^*2*^ = 99%) and largely due to variation between studies (*I*^*2*^ = 91%), with only a small amount of residual variation (*I*^*2*^ = 8%). There was no evidence of changes in effect sizes across time (*β*_*year*_ = 0.0036, 95%CI = −0.0012 – 0.0084, z = 1.46, P = 0.145). Finally, there was no evidence of publication bias through inspection of funnel plots (Figure S3) or regression of model residuals on effect size variances in either the full data set (*t* = −0.30, DF = 74, P = 0.763) or published data (*t* = −0.03, DF = 14, P = 0.975).

**Table 3:**
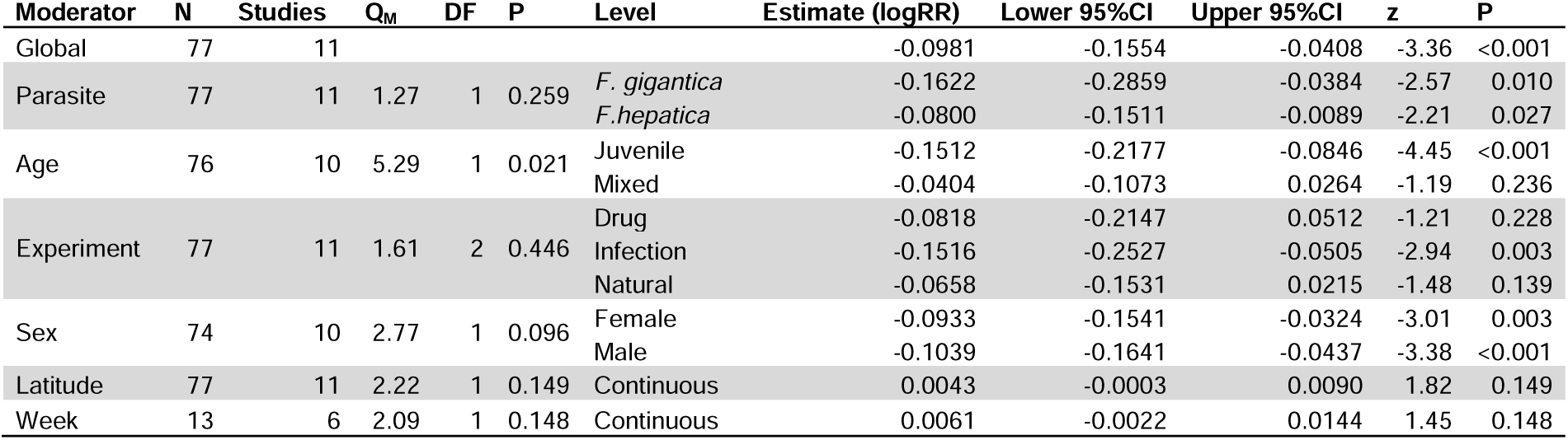
Results from meta-analysis of daily weight gain (DWG). ‘N’ is the number of effects sizes available for analysis. ‘Global’ indicates the overall effect size from the initial random effects model, while the subsequent moderator effects are from mixed-effects models where each moderator was fitted in turn. ‘Effect sizes’ and ‘Studies’ varies because information was not available for some of the effect sizes. ‘Q_M_’ and the associated ‘DF’ and ‘P’ values refer to the Wald-type test for significant differences in effect size due to moderator, while ‘z’ and ‘P’ values refer to tests for differences from zero for different levels of each moderator.

| Moderator | N | Studies | Q<sub>M</sub> | DF | P | Level | Estimate (logRR) | Lower 95%CI | Upper 95%CI | z | P |
| --- | --- | --- | --- | --- | --- | --- | --- | --- | --- | --- | --- |
| Global | 77 | 11 |  |  |  |  | -0.0981 | -0.1554 | -0.0408 | -3.36 | <0.001 |
| Parasite | 77 | 11 | 1.27 | 1 | 0.259 | *F. gigantica* | -0.1622 | -0.2859 | -0.0384 | -2.57 | 0.010 |
|  |  |  |  |  |  | *F. hepatica* | -0.0800 | -0.1511 | -0.0089 | -2.21 | 0.027 |
| Age | 76 | 10 | 5.29 | 1 | 0.021 | Juvenile | -0.1512 | -0.2177 | -0.0846 | -4.45 | <0.001 |
|  |  |  |  |  |  | Mixed | -0.0404 | -0.1073 | 0.0264 | -1.19 | 0.236 |
| Experiment | 77 | 11 | 1.61 | 2 | 0.446 | Drug | -0.0818 | -0.2147 | 0.0512 | -1.21 | 0.228 |
|  |  |  |  |  |  | Infection | -0.1516 | -0.2527 | -0.0505 | -2.94 | 0.003 |
|  |  |  |  |  |  | Natural | -0.0658 | -0.1531 | 0.0215 | -1.48 | 0.139 |
| Sex | 74 | 10 | 2.77 | 1 | 0.096 | Female | -0.0933 | -0.1541 | -0.0324 | -3.01 | 0.003 |
|  |  |  |  |  |  | Male | -0.1039 | -0.1641 | -0.0437 | -3.38 | <0.001 |
| Latitude | 77 | 11 | 2.22 | 1 | 0.149 | Continuous | 0.0043 | -0.0003 | 0.0090 | 1.82 | 0.149 |
| Week | 13 | 6 | 2.09 | 1 | 0.148 | Continuous | 0.0061 | -0.0022 | 0.0144 | 1.45 | 0.148 |

**Figure 1.**
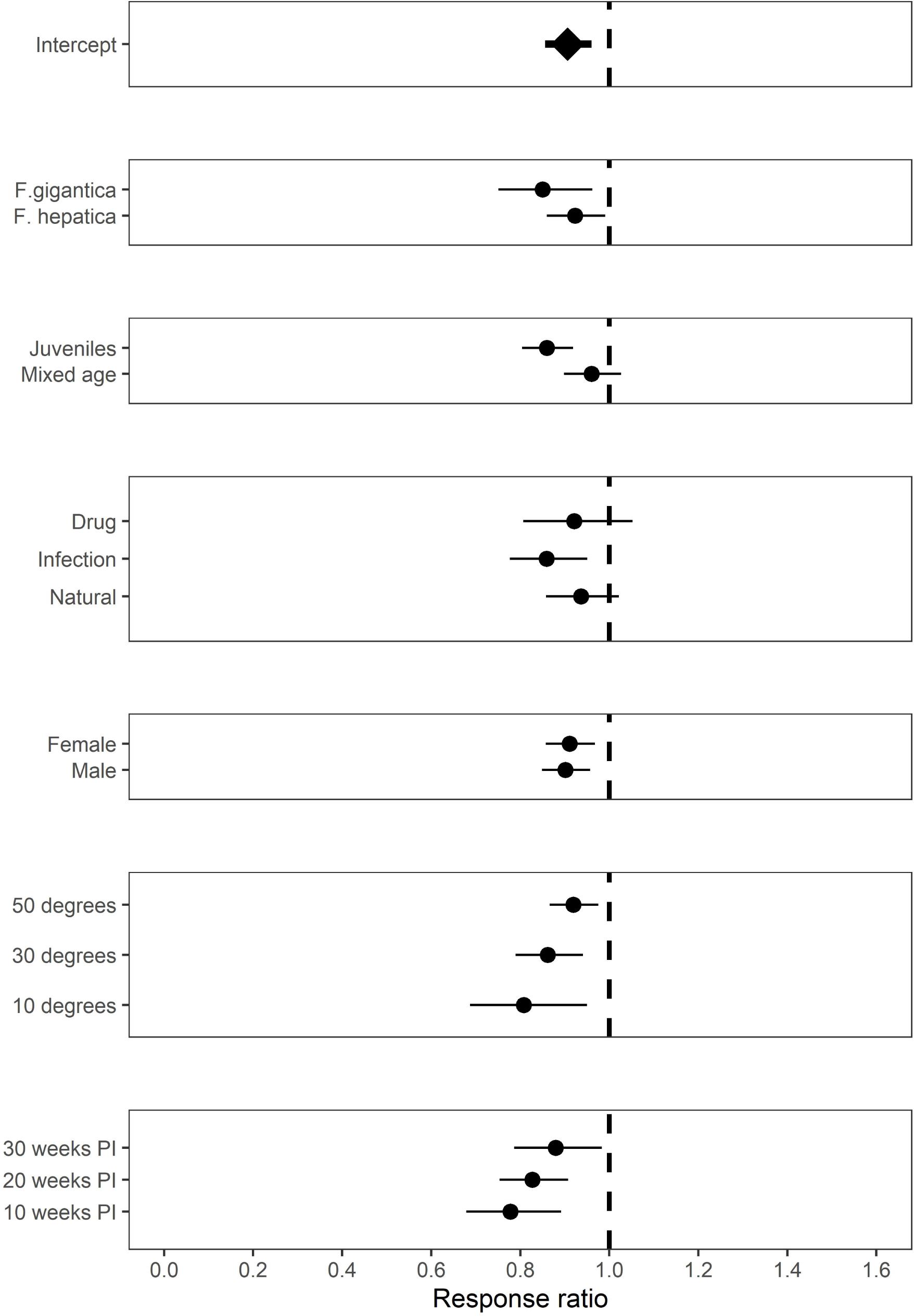
Estimated response ratio (RR) from meta-regression analysis of fluke-infected and uninfected animals for daily weight gain (DWG). A response ratio of 1 indicates equal performance in infected and uninfected animals; values <1 indicate poorer performance in infected animals. Plots show estimated mean effect of the global effect on infection (Intercept), plus the effect of infection in different moderators, with 95%CI.

### Live weight

Multi-level meta-analysis of live weight revealed a negative effect of fluke infection (*β*_*global*_ = – 0.0611, 95%CI = −0.0877 – −0.0344, z = −4.48, P < 0.001), suggesting that infected animals weighed around 6% less than uninfected animals (Figure 2). Meta-regression revealed that the moderators of parasite species, host species, host age, experimental type, and latitude had no influence on the effect size (Table 4). There was, however, a significant effect of sex (Q_M_ = 16.04, P < 0.001), with males and especially mixed-sex groups of animals showing a greater impact of fluke than female only groups (Figure 2). There was also support for week post-infection (Q_M_ = 7.66, P = 0.006), with effects of fluke increasing across time in experimental fluke challenge studies (Figure S4). Fluke had a negative impact on live weight in animals infected with *F. hepatica* and *F. gigantica* (although the latter effect was marginally not supported), in both cattle and sheep, in young animals but not adults, in studies that administered experimental infections rather than those that cleared infections with flukicides and in male and mixed-sex groups but not females (Table 4). Total variation in effect sizes was moderate (*I*^*2*^ = 71%) and largely due to residual variation (*I*^*2*^ = 50%), with smaller variation due to between-study effects (*I*^*2*^ = 22%). Effect sizes did not change with time (*β*_*year*_ = −0.0006, 95%CI = −0.0023 – 0.0012, z = −0.61, P = 0.542). Finally, there was no evidence of publication bias through inspection of funnel plots (Figure S5) or regression of model residuals on effect size variances (*t* = −0.41, P = 0.683).

**Table 4.** Results from meta-analysis of live weight. ‘N’ is the number of effects sizes available for analysis. ‘Global’ indicates the overall effect size from the initial random effects model, while the subsequent moderator effects are from mixed-effects models where each moderator was fitted in turn. ‘Effect sizes’ and ‘Studies’ varies because information was not available for some of the effect sizes. ‘Q_M_’ and the associated ‘DF’ and ‘P’ values refer to the Wald-type test for significant differences in effect size due to moderator, while ‘z’ and ‘P’ values refer to tests for differences from zero for different levels of each moderator.

| Moderator | N | Studies | Q<sub>M</sub> | DF | P | Level | Estimate (logRR) | Lower 95%CI | Upper 95%CI | z | P |
| --- | --- | --- | --- | --- | --- | --- | --- | --- | --- | --- | --- |
| Global | 47 | 11 |  |  |  |  | -0.0611 | -0.0877 | -0.0344 | -4.48 | <0.001 |
| Parasite | 46 | 10 | 0.01 | 1 | 0.913 | *F. gigantica* | -0.0519 | -0.1109 | 0.0070 | -1.73 | 0.084 |
|  |  |  |  |  |  | *F. hepatica* | -0.0556 | -0.0860 | -0.0253 | -3.59 | 0.003 |
| Host | 47 | 11 | 0.01 | 1 | 0.910 | Cattle | -0.0598 | -0.0960 | -0.0236 | -3.24 | 0.001 |
|  |  |  |  |  |  | Sheep | -0.0631 | -0.1076 | -0.0186 | -2.78 | 0.005 |
| Age | 47 | 11 | 2.38 | 1 | 0.123 | Adult | -0.0222 | -0.0757 | 0.0312 | -0.81 | 0.415 |
|  |  |  |  |  |  | Juvenile | -0.0704 | -0.0973 | -0.0435 | -5.12 | <0.001 |
| Experiment | 46 | 10 | 1.84 | 1 | 0.174 | Drug | -0.0225 | -0.0743 | 0.0293 | -0.85 | 0.395 |
|  |  |  |  |  |  | Infection | -0.0642 | -0.0915 | -0.0368 | -4.60 | <0.001 |
| Sex | 47 | 11 | 16.04 | 2 | <0.001 | Female | -0.0234 | -0.0518 | 0.0050 | -1.62 | 0.106 |
|  |  |  |  |  |  | Male | -0.0478 | -0.0785 | -0.0172 | -3.06 | 0.002 |
|  |  |  |  |  |  | Mixed | -0.1042 | -0.1330 | -0.0755 | -7.10 | <0.001 |
| Latitude | 47 | 11 | 0.15 | 1 | 0.696 | Continuous | -0.0003 | -0.0021 | 0.0014 | -0.39 | 0.696 |
| Week | 33 | 8 | 7.66 | 1 | 0.006 | Continuous | -0.0023 | -0.0039 | -0.0007 | -2.77 | 0.006 |

**Figure 2.**
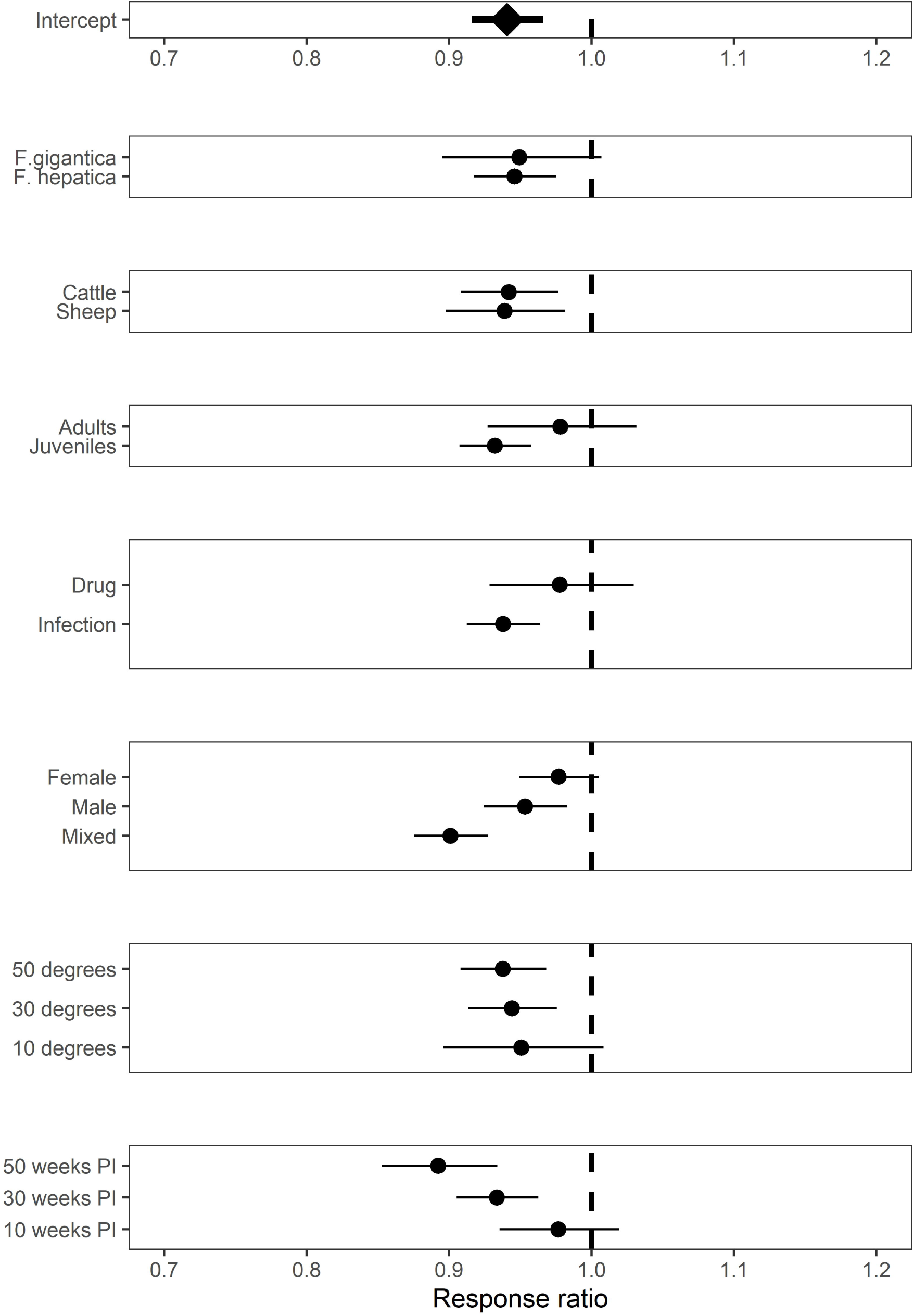
Estimated response ratio (RR) from meta-regression analysis of fluke-infected and uninfected animals for live weight. A response ratio of 1 indicates equal performance in infected and uninfected animals; values <1 indicate poorer performance in infected animals. Plots show estimated mean effect of the global effect on infection (Intercept), plus the effect of infection in different moderators, with 95%CI.

### Carcass weight

Analysis of carcass weight supported a significant negative effect of fluke infection (*β*_*global*_ = – 0.0060, 95%CI = −0.0099 – −0.0021, z = −3.00, P = 0.003), although the effect size was negligible, suggesting a 0.6% reduction in carcass weight in infected animals (Figure 3).

**Figure 3.**
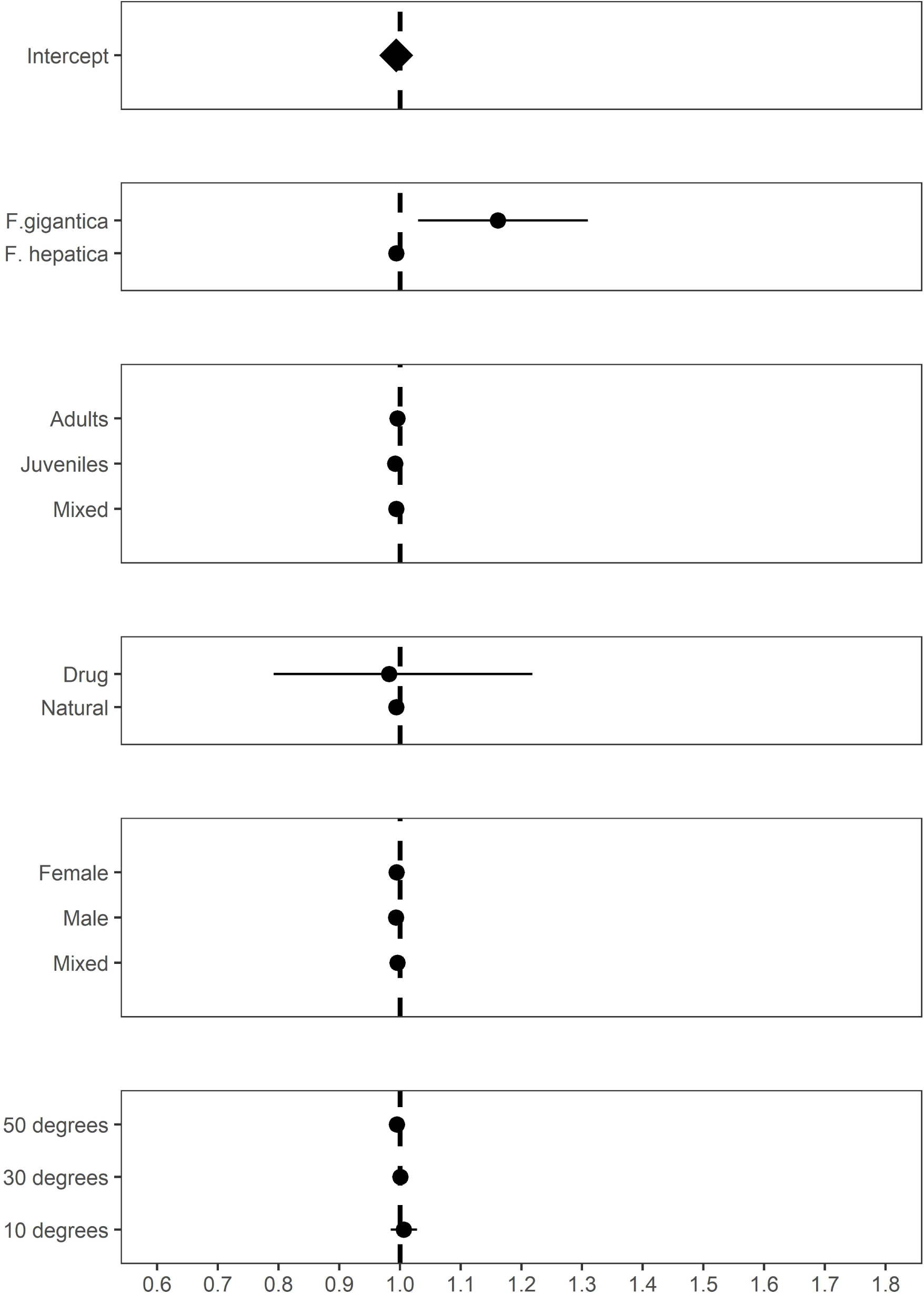
Estimated response ratio (RR) from meta-regression analysis of fluke-infected and uninfected animals for carcass weight. A response ratio of 1 indicates equal performance in infected and uninfected animals; values <1 indicate poorer performance in infected animals. Plots show estimated mean effect of the global effect on infection (Intercept), plus the effect of infection in different moderators, with 95%CI.

Meta-regression revealed that the moderators of age, experimental type, and latitude had no influence on the effect size (Table 5). There was, however, a significant effect of parasite species (Q_M_ = 6.44, P = 0.011), with animals infected with *F. gigantica* having greater carcass weight than uninfected animals, and animals infected with *F. hepatica* having marginally lower carcass weight than uninfected animals (Figure 3). Total variation in effect sizes was high (*I*^*2*^ = 82%), with a relatively small influence of between-study effects (23%) and a large contribution of residual variation (59%). Effect sizes increased significantly with time (*β*_*year*_ = 0.0013, 95%CI = 0.0005 – 0.0021, z = 3.21, P = 0.001). There was no evidence of publication bias through inspection of funnel plots (Figure S6) or regression analysis in either the full dataset (*t* = 0.06, DF = 82, P = 0.951) or the published data (*t* = 0.39, DF = 22, P = 0.701).

**Table 5.** Results from meta-analysis of carcass weight. ‘N’ is the number of effects sizes available for analysis. ‘Global’ indicates the overall effect size from the initial random effects model, while the subsequent moderator effects are from mixed-effects models where each moderator was fitted in turn. ‘Effect sizes’ and ‘Studies’ varies because information was not available for some of the effect sizes. ‘Q_M_’ and the associated ‘DF’ and ‘P’ values refer to the Wald-type test for significant differences in effect size due to moderator, while ‘z’ and ‘P’ values refer to tests for differences from zero for different levels of each moderator.

| Moderator | N | Studies | Q<sub>M</sub> | DF | P | Level | Estimate (logRR) | Lower 95%CI | Upper 95%CI | z | P |
| --- | --- | --- | --- | --- | --- | --- | --- | --- | --- | --- | --- |
| Global | 84 | 8 |  |  |  |  | -0.0060 | -0.0099 | -0.0021 | -3.00 | 0.003 |
| Parasite | 84 | 8 | 6.44 | 1 | 0.011 | *F. gigantica* | 0.1497 | 0.0294 | 0.2700 | 2.44 | 0.015 |
|  |  |  |  |  |  | *F. hepatica* | -0.0062 | -0.0101 | 0.0023 | -3.10 | 0.002 |
| Age | 84 | 8 | 0.37 | 2 | 0.831 | Adult | -0.0042 | -0.0127 | 0.0044 | -0.96 | 0.340 |
|  |  |  |  |  |  | Juvenile | -0.0081 | -0.0209 | 0.0048 | -1.23 | 0.219 |
|  |  |  |  |  |  | Mixed | -0.0063 | -0.0114 | -0.0011 | -2.40 | 0.017 |
| Experiment | 84 | 8 | 0.01 | 1 | 0.914 | Drug | -0.0178 | -0.2332 | 0.1975 | -0.16 | 0.871 |
|  |  |  |  |  |  | Natural | -0.0060 | -0.0099 | -0.0021 | -2.99 | 0.003 |
| Sex | 81 | 7 | 0.32 | 2 | 0.851 | Female | -0.0056 | -0.0102 | -0.0009 | -2.34 | 0.019 |
|  |  |  |  |  |  | Male | -0.0066 | -0.0113 | -0.0019 | -2.78 | 0.006 |
|  |  |  |  |  |  | Mixed | -0.0043 | -0.0170 | 0.0084 | -0.67 | 0.506 |
| Latitude | 84 | 8 | 1.30 | 1 | 0.254 | Continuous | -0.0003 | -0.0008 | 0.0002 | -1.13 | 0.257 |

### Total weight gain

There was an overall negative influence of fluke infection on total weight gain, and while the effect did not reach statistical significance, it was of considerable magnitude (*β*_*global*_ = – 0.1541, 95%CI = −0.3258 – 0.0176, z = −1.76, P = 0.079), with infected animals gaining 14% less weight than infected animals (Figure S7). Meta-regression revealed that the moderators of parasite species, experimental type, sex, latitude and week had no influence on the effect size (Table S1). Negative effects of fluke were detected in studies that administered experimental infections rather than drug treatments, and in males but not mixed-sex groups (Figure S7). Total variation in effect sizes was high (*I*^*2*^ = 79%) and largely due to variation between breeds (*I*^*2*^ = 72%), with smaller amounts of between-study (*I*^*2*^ = 4%) and residual variation (*I*^*2*^ = 2%). There was no evidence of changes in effect sizes across time (*β*_*year*_ = 0.0040, 95%CI = −0.0086 – 0.0166, z = 0.62, P = 0.537) and no evidence of publication bias through funnel plots (Figure S8) or regression analysis (*t* = 0.85, P = 0.409).

### Milk production

There was no support for a significant overall influence of fluke infection on milk production (*β*_*global*_ = −0.0500, 95%CI = −0.1065 – 0.0066, z = −1.73, P = 0.083; Figure S9). Meta-regression revealed that the moderators of experimental type and latitude had no influence on the effect size (Table S2). Total variation in effect sizes was high (*I*^*2*^ = 88%), but we did not fit random effects of study or breed because they were found to be zero. There was no evidence of changes in effect sizes across time (*β*_*year*_ = −0.0004, 95%CI = −0.0040 – 0.0031, z = −0.23, P = 0.815) and no evidence of publication bias through inspection of a funnel plot (Figure S10) and regression analysis (*t* = −0.17, P = 0.875).

## DISCUSSION

In this study, we used a meta-analytic approach in order to estimate the overall influence of liver fluke infection on performance traits in sheep and cattle. We found large amounts of variation between studies in the effects they estimated for liver fluke in each trait, and identified moderator variables that were associated with effect sizes. For each trait, we initially ran a random-effects meta-analysis in order to estimate an overall effect size. We found significant and substantial effects of fluke infection on two traits, with a 9% reduction in daily weight gain (DWG) and a 6% reduction in live weight. Moreover, we found a significant reduction in carcass weight in fluke-infected animals, although the overall effect was negligible, with infected animals having 0.6% lower carcass weight than fluke-free animals. Reductions in total weight gain and milk production in fluke-infected animals were not statistically supported. We also found that, although moderators were generally not supported, there was evidence to suggest that studies were more likely to find significant effects of fluke if they used experimental infection compared to natural infections or infected versus flukicide-treated animals, and if they studied young animals rather than adults.

The overall effect of a reduced DWG of 9% is likely to have a significant impact on several parameters: for the welfare of the animal, experiencing chronic disease (Howell and Williams, 2020); for the producer, in terms of financial costs (Mehmood et al., 2017; Schweizer et al., 2005); and for the environment, with the increased greenhouse gas emissions emitted by less efficient animals (Skuce et al., in prep). There was support for the moderator of age group: the effect of fluke was significantly greater in young animals compared to mixed-age animals. This is potentially related to the fact that young animals are likely to be growing, while older animals may be putting on weight rather than growing *per se*; as such, younger animals are more likely to suffer the effects of fluke infection with regard to their ability to increase in weight. Further, it is possible that this may reflect age-dependent effects of immunity to the parasite, with younger animals less able to control fluke infection – although evidence for effective immunity in adults is scant (Hoyle et al., 2003) – or responding in a way which induces more immunopathology. While the moderator of experimental design was not supported, infection studies showed an effect of fluke that was significantly different from zero (Table 3), but this was not the case for drug studies or studies observing the effects of natural infections. Infection studies may be associated with larger effect sizes because they tended to involve administration of a single large infective dose of metacercariae at the start of the experiment; of the 15 experimental infection studies across our whole dataset, only two used trickle infections. It may well be that a large influx of immature fluke caused more damage that a natural infection due to the migration of a large number of larvae simultaneously (Boray, 1967; Ross and Dow, 1966). Another explanation is that we may have seen smaller effects in drug studies because they may not have fully cleared fluke infections (even though mean burdens in treated animals were very low), and in natural infection studies because unlike in experimental challenge studies, there would be variation in fluke challenge and duration of infection within infected groups resulting in a more variable response to infection. Finally, we saw significant effects of both *F. hepatica* and *F. gigantica* on DWG, and very similar effects in both sexes. There was, however, no variation in effect size with latitude or across time in experimental studies.

The difference in live weight between infected and uninfected animals of 6% is, as with DWG, likely to prove relevant to the efficiency of production. Two moderators were supported. First, there was variation between sex classes, with stronger effects of fluke in males and mixed-sex groups than in females. This is likely to be due to the fact that all effect sizes of males and mixed-sex groups came from studies on young animals, while most of those on females (12/14) came from adults, and as with DWG, there were stronger effects of fluke in young animals. There was also a significant effect of duration of infection, with effect sizes increasing with week post-infection in experimental infection studies; this result is consistent with our finding of lower DWG in infected animals, since a difference in DWG would lead to an increasing divergence in live weight across time. As with DWG, there were stronger effects of fluke in young animals compared to adults, and in infection experiments compared to drug experiments, presumably for the same reasons as discussed above for DWG. A final striking observation was the very similar magnitude of the average effect size in cattle and sheep, which is potentially surprising given the generally greater capacity for fluke to cause more severe disease in sheep than cattle (Howell and Williams, 2020).

While we found a significant impact of fluke infection on carcass weight, the overall effect was negligible, with infected animals having carcass weights only 0.6% lower than uninfected animals. The statistical significant of such a small effect size is likely related to the fact that the majority of the data came from abattoir studies, with very large sample sizes and very small effect size variances. The utility of carcass weight as a metric for assessing performance may be questionable, given that most producers will take animals to slaughter only when they have reached a target weight – and indeed will be penalised for not doing so – yet carcass weight is still commonly reported as a measure of performance. The only noteworthy moderator variable was that of fluke species: animals with *F. gigantica* actually had higher carcass weights than uninfected animals. That said, all of the effect sizes concerning *F. gigantica* came from a single study (Molina, 2005), in which three groups of cattle of different ages were heavier when infected with *F. gigantica* compared to their uninfected counterparts. In the same abattoir study, water buffalo (*Bubalus bubalis*, which were not included in the meta-analysis) had lighter carcasses when infected with fluke, and no explanation was apparent or discussed (Molina, 2005). The overall effect size for total weight gain was substantial, with 14% lower total weight gain in fluke-infected animals, although this was marginally non-significant. Once again, we saw a considerably stronger effect in infection experiments compared to drug studies.

While traits relating to body weight were well represented, we had a much smaller dataset for milk yield. Many studies have tested the impact of fluke infection on milk yield and have found substantial effects (Arenal et al., 2018; Charlier et al., 2007; Howell et al., 2015; Köstenberger et al., 2017; Mezo et al., 2011), but these have studied effects at the herd level using bulk tank milk samples and so were not included in our analysis. Other studies of individual animals did not report the necessary statistics for us to calculate effect size variance (Khan et al., 2010). Perhaps due to the small number of studies included in our analyses, we found no overall influence for an effect of fluke on milk yield. A future meta-analysis of the many bulk tank studies could prove informative in determining overall herd-level effects.

The effect of fluke infection has been measured with respect to other performance traits, as reviewed recently (Charlier et al., 2013; Skuce and Zadoks, 2013), but for these there were insufficient studies in our initial scoping search for them to be included in the full search and analysis. For example, there have been many studies of the influence of fluke infection on wool weight, but during our initial searches, we found that most of these did not report the standard deviation (or standard errors and sample sizes) that were required to calculate effect size variance. These older studies did typically report substantial effects of fluke infection on wool yield: two studies administering a range of infection doses reported lower yields in infected animals, particularly at higher infectious doses (Edwards et al., 1976; Hawkins and Morris, 1978) and a study testing the effect of flukicide treatment found that treatment enhanced wool growth (Hawkins, 1984). Two further studies reported increasing effects of fluke infection on wool yield over time (Hagh-Nazari and Dalimi, 2000; Roseby, 1970), suggesting that larger effects are seen once adult fluke are established in the liver from 11-12 weeks post-infection (Kaplan, 2001). In addition, several studies report a negative influence of fluke on reproduction (Marley et al., 1994), including a 39 day delay in first oestrus in cattle (López-Díaz et al., 1998), a 4.7 day increase in calving interval (Charlier et al., 2007) and an increase in pregnancy rate from 57% in control ewes to 80% in flukicide-treated ewes (Hope Cawdery, 1976). Some studies, however, report no significant influence of fluke infection on reproductive traits (Loyacano et al., 2002; Mezo et al., 2011). Another performance trait that we did not consider was time to reach slaughter weight, which has been found to be delayed by 33-93 days in fluke-infected animals, depending on the severity of infection (Mazeri et al., 2017), but once again, there were too few studies to consider this as an outcome trait.

Only a small number of meta-analyses have provided quantitative reviews of the impacts of disease on performance in livestock and in general have reported stronger effects than we saw here. A meta-analysis of 75 trials studying the impact of anthelmintic treatment on milk production in dairy cows reported an increased daily yield of 0.35kg in treated compared to untreated animals, suggesting a negative influence of helminth infection (almost exclusively gastrointestinal nematodes based on the studies included) on performance (Sanchez et al., 2004). A meta-analysis of 18 studies examining the influence of endoparasite – largely nematode – infection on weight gain in pigs reported a 31% reduction in infected animals, which was largely due to reduced feed intake in infected animals (Kipper et al., 2011). More closely related to our study, a meta-analysis of 94 effect sizes examining effects of nematode infections on sheep performance across Europe found that nematode infection was associated with a 15% reduction in weight gain, 10% lower wool production and 22% lower milk yield (Mavrot et al., 2015). While this latter study was unable to estimate the impact of moderator variables on outcomes, it was able to show that increased parasite burden, measured as faecal egg count, was associated with a greater negative impact of infection in “highly parasitized” compared to “low parasitized” individuals (Mavrot et al., 2015). We were unable to estimate the impact of fluke burden as a moderator in our study because of the large variation between studies in their design: in most studies, individuals were simply assigned as “infected” or “uninfected”, and in experimental infection studies, a range of infection doses were given, either as single infections or as trickles across varying time scales.

## CONCLUSION

This meta-analysis has provided quantitative estimates of the impact of liver fluke infection on the performance of sheep and cattle by collating relevant data from both published and unpublished sources. We found that effects on live weight gain and live weight were the most pronounced, while effects on carcass weight and milk production were either negligible or non-significant. Since we focused on individual-based assessments of infection, there remains a great deal of data to be exploited for meta-analysis of influences of fluke at the herd level, particularly regarding bulk-tank milk samples. Our results also reveal that studies administering experimental infections, and studies of younger animals, are more likely to reveal effects of fluke that differ from zero, and that detecting effects of natural infection may be more difficult. Improved diagnostics of fluke infection, particularly those which can quantify the level of infection, may be required to gain a more accurate estimate of the impact of subclinical fluke infection on performance in the future, aiding understanding of between-host variation in resistance and tolerance of infection and aiding the effort to design improved mitigation strategies for fluke infection.

## ACKNOWLEDGEMENTS

We are deeply indebted to those people who took the trouble to provide us with data from their published studies: Camille Bellet, Ricardo Almeida da Costa, Manuel Sanchez-Vazquez and Alan Twomey. We are also grateful to Harbro Ltd, Innovent Technology Ltd, McIntosh-Donald Ltd, Scotbeef Ltd, Willie Thomson and David Barclay for allowing us to include some of their data in the analysis. ADH is funded by a Moredun Foundation Fellowship.

## AUTHOR CREDIT STATEMENT

**Adam Hayward:** conceptualization, formal analysis, investigation, data curation, writing – original draft, writing – review and editing, visualization

**Philip Skuce:** writing – review and editing; supervision

**Tom McNeilly:** writing – review and editing, supervision

## REFERENCES

Arenal, A., García, Y., Quesada, L., Velázquez, D., Sánchez, D., Peña, M., Suárez, A., Díaz, A., Sánchez, Y., Casaert, S., van Dijk, J., Vercruysse, J., Charlier, J., 2018. Risk factors for the presence of Fasciola hepatica antibodies in bulk-milk samples and their association with milk production decreases, in Cuban dairy cattle. BMC Veterinary Research 14, 336.

Beesley, N.J., Caminade, C., Charlier, J., Flynn, R.J., Hodgkinson, J.E., Martinez-Moreno, A., Martinez-Valladares, M., Perez, J., Rinaldi, L., Williams, D.J.L., 2018. Fasciola and fasciolosis in ruminants in Europe: Identifying research needs. Transboundary and Emerging Diseases 65, 199–216.

Bellet, C., Green, M.J., Vickers, M., Forbes, A., Berry, E., Kaler, J., 2016. Ostertagia spp., rumen fluke and liver fluke single- and poly-infections in cattle: An abattoir study of prevalence and production impacts in England and Wales. Preventive Veterinary Medicine 132, 98–106.

Berry, C.I., 1977. Studies on the pathogenesis of ovine fascioliasis and schistosomiasis. University of Glasgow, Glasgow.

Boray, J.C., 1967. Studies on experimental infections with Fasciola hepatica, with particular reference to acute fascioliasis in sheep. Annals of Tropical Medicine & Parasitology 61, 439–450.

Bossaert, K., Farnir, F., Leclipteux, T., Protz, M., Lonneux, J.-F., Losson, B., 2000. Humoral immune response in calves to single-dose, trickle and challenge infections with Fasciola hepatica. Veterinary Parasitology 87, 103–123.

Brockwell, Y.M., Elliott, T.P., Anderson, G.R., Stanton, R., Spithill, T.W., Sangster, N.C., 2013. Confirmation of Fasciola hepatica resistant to triclabendazole in naturally infected Australian beef and dairy cattle. Int J Parasitol Drugs Drug Resist 4, 48–54.

Charlier, J., De Cat, A., Forbes, A., Vercruysse, J., 2009. Measurement of antibodies to gastrointestinal nematodes and liver fluke in meat juice of beef cattle and associations with carcass parameters. Veterinary Parasitology 166, 235–240.

Charlier, J., Duchateau, L., Claerebout, E., Williams, D., Vercruysse, J., 2007. Associations between anti-Fasciola hepatica antibody levels in bulk-tank milk samples and production parameters in dairy herds. Preventive Veterinary Medicine 78, 57–66.

Charlier, J., Vercruysse, J., Morgan, E., Van Dijk, J., Williams, D.J.L., 2013. Recent advances in the diagnosis, impact on production and prediction of Fasciola hepatica in cattle. Parasitology 141, 326–335.

Chick, B.F., Coverdale, O.R., Jackson, A.R.B., 1980. Production effects of liver fluke (Fasciola hepatica) infection in beef cattle. Australian Veterinary Journal 56, 588–592.

Crossland, N.O., Johnstone, A., Beaumont, G., Bennett, M.S., 1977. The Effect of Control of Chronic Fascioliasis on the Productivity of Lowland Sheep. British Veterinary Journal 133, 518–525.

da Costa, R.A., Corbellini, L.G., Castro-Janer, E., Riet-Correa, F., 2019. Evaluation of losses in carcasses of cattle naturally infected with Fasciola hepatica: effects on weight by age range and on carcass quality parameters. International Journal for Parasitology 49, 867–872.

Dargie, J.D., 1980. The pathophysiological effects of gastrointestinal and liver parasites in sheep, In: Ruckebusch, Y., Thivend, P. (Eds.) Digestive Physiology and Metabolism in Ruminants: Proceedings of the 5th International Symposium on Ruminant Physiology, held at Clermont — Ferrand, on 3rd–7th September, 1979. Springer Netherlands, Dordrecht, pp. 349–371.

Dargie, J.D., 1987. The impact on production and mechanisms of pathogenesis of trematode infections in cattle and sheep. International Journal for Parasitology 17, 453–463.

Echevarria, F.A.M., Correa, M.B.C., Wehrle, R.D., Correa, I.F., 1992. Experiments on anthelmintic control of Fasciola hepatica in Brazil. Veterinary Parasitology 43, 211–222.

Edwards, C.M., al-Saigh, M.N., Williams, G.L., Chamberlain, A.G., 1976. Effect of liver fluke on wool production in Welsh mountain sheep. Vet Rec 98, 372.

El-Tahawy, A.S., Bazh, E.K., Khalafalla, R.E., 2017. Epidemiology of bovine fascioliasis in the Nile Delta region of Egypt: Its prevalence, evaluation of risk factors, and its economic significance. Vet World 10, 1241–1249.

Elelu, N., Eisler, M.C., 2018. A review of bovine fasciolosis and other trematode infections in Nigeria. Journal of Helminthology 92, 128–141.

Forbes, A.B., Reddick, D., Stear, M.J., 2015. Efficacy of treatment of cattle for liver fluke at housing: influence of differences in flukicidal activity against juvenile Fasciola hepatica. Veterinary Record 176, 333–333.

French, A.S., Zadoks, R.N., Skuce, P.J., Mitchell, G., Gordon-Gibbs, D.K., Taggart, M.A., 2019. Habitat and host factors associated with liver fluke (Fasciola hepatica) diagnoses in wild red deer (Cervus elaphus) in the Scottish Highlands. Parasites & Vectors 12, 535.

Genicot, B., Mouligneau, F., Lekeux, P., 1991. Economic and production consequences of liver fluke disease in double-muscled fattening cattle. Journal of Veterinary Medicine, Series B 38, 203–208.

Glass, G.V., 1976. Primary, secondary, and meta-analysis of research. Educational Researcher 5, 3–8.

Gurevitch, J., Koricheva, J., Nakagawa, S., Stewart, G., 2018. Meta-analysis and the science of research synthesis. Nature 555, 175–182.

Hagh-Nazari, J., Dalimi, A., 2000. Effect of Fasciola gigantica infection on the quality and quantity of wool production of Baluchi sheep. The Indian Journal of Animal Sciences 70, 271–273.

Hawkins, C.D., 1984. Productivity in sheep treated with diamphenethide at different times after infection with Fasciola hepatica. Veterinary Parasitology 15, 117–123.

Hawkins, C.D., Morris, R.S., 1978. Depression of productivity in sheep infected with Fasciola hepatica. Veterinary Parasitology 4, 341–351.

Hope Cawdery, M.J., 1976. The effects of fascioliasis on ewe fertility. British Veterinary Journal 132, 568–575.

Hope Cawdery, M.J., Strickland, K.L., Conway, A., Crowe, P.J., 1977. Production effects of liver fluke in cattle I. the effects of infection on liveweight gain, feed intake and food conversion efficiency in beef cattle. British Veterinary Journal 133, 145–159.

Howell, A., Baylis, M., Smith, R., Pinchbeck, G., Williams, D., 2015. Epidemiology and impact of Fasciola hepatica exposure in high-yielding dairy herds. Preventive Veterinary Medicine 121, 41–48.

Howell, A.K., Williams, D.J.L., 2020. The epidemiology and control of liver flukes in cattle and sheep. Veterinary Clinics of North America: Food Animal Practice 36, 109–123.

Hoyle, D.V., Dalton, J.P., Chase-Topping, M., Taylor, D.W., 2003. Pre-exposure of cattle to drug-abbreviated Fasciola hepatica infections: the effect upon subsequent challenge infection and the early immune response. Veterinary Parasitology 111, 65–82.

Jacob, A., Singh, P., Verma, A.K., 2014. Effect of supplementation of deoiled mahua seed cake on the growth performance and blood biochemical parameters of crossbred calves during recovery period of infection from F. gigantica. Animal Nutrition and Feed Technology 14, 161–168.

Jacob, A.B., Singh, P., Verma, A.K., 2015. Effect of feeding deoiled mahua (Bassia latifolia) seed cake on the growth performance, digestibility and balance of nutrients in cross-bred calves during pre-patent period of Fasciola gigantica infection. Journal of Animal Physiology and Animal Nutrition 99, 299–307.

Kamaludeen, J., Graham-Brown, J., Stephens, N., Miller, J., Howell, A., Beesley, N.J., Hodgkinson, J., Learmount, J., Williams, D., 2019. Lack of efficacy of triclabendazole against Fasciola hepatica is present on sheep farms in three regions of England, and Wales. Veterinary Record 184, 502–502.

Kaplan, R.M., 2001. Fasciola hepatica: a review of the conomic impact in cattle and considerations for control. Veterinary Therapeutics 2, 40–50.

Khan, M.K., Sajid, M.S., Khan, M.N., Iqbal, Z., Arshad, M., Hussain, A., 2010. Point prevalence of bovine fascioliasis and the influence of chemotherapy on the milk yield in a lactating bovine population from the district of Toba Tek Singh, Pakistan. Journal of Helminthology 85, 334–338.

Kipper, M., Andretta, I., Monteiro, S.G., Lovatto, P.A., Lehnen, C.R., 2011. Meta-analysis of the effects of endoparasites on pig performance. Veterinary Parasitology 181, 316–320.

Köstenberger, K., Tichy, A., Bauer, K., Pless, P., Wittek, T., 2017. Associations between fasciolosis and milk production, and the impact of anthelmintic treatment in dairy herds. Parasitology Research 116, 1981–1987.

Lau, J., Rothstein, H.R., Stewart, G.B., 2013. History and progress of meta-analysis, In: Koricheva, J., Gurevitch, J., Mengersen, K. (Eds.) Handbook of Meta-Analysis in Ecology and Evolution. Princeton University Press, Princeton, pp. 407–419.

Lean, I.J., Rabiee, A.R., Duffield, T.F., Dohoo, I.R., 2009. Use of meta-analysis in animal health and reproduction: Methods and applications. Journal of Dairy Science 92, 3545–3565.

López-Abán, J., Casanueva, P., Nogal, J., Arias, M., Morrondo, P., Diez-Baños, P., Hillyer, G.V., Martínez-Fernández, A.R., Muro, A., 2007. Progress in the development of Fasciola hepatica vaccine using recombinant fatty acid binding protein with the adjuvant adaptation system ADAD. Veterinary Parasitology 145, 287–296.

López-Díaz, M.C., Carro, M.C., Cadórniga, C., Díez-Baños, P., Mezo, M., 1998. Puberty and serum concentrations of ovarian steroids during prepuberal period in friesian heifers artificially infected with Fasciola hepatica. Theriogenology 50, 587–593.

Loyacano, A.F., Williams, J.C., Gurie, J., DeRosa, A.A., 2002. Effect of gastrointestinal nematode and liver fluke infections on weight gain and reproductive performance of beef heifers. Veterinary Parasitology 107, 227–234.

Mage, C., Levieux, D., Bernabe, P., Degez, P., 1993. Liver fluke therapy by closantel in culled dairy cows. Revue de Medecine Veterinaire (France).

Marley, S., Knapp, S., Johnson, G., 1994. Performance of calves from liver fluke infected heifers treated in western Montana. Agri-Practice (USA).

Mavrot, F., Hertzberg, H., Torgerson, P., 2015. Effect of gastro-intestinal nematode infection on sheep performance: a systematic review and meta-analysis. Parasites & Vectors 8, 557.

May, K., Bohlsen, E., König, S., Strube, C., 2020. Fasciola hepatica seroprevalence in Northern German dairy herds and associations with milk production parameters and milk ketone bodies. Veterinary Parasitology 277, 109016.

May, K., Brügemann, K., König, S., Strube, C., 2019. Patent infections with Fasciola hepatica and paramphistomes (Calicophoron daubneyi) in dairy cows and association of fasciolosis with individual milk production and fertility parameters. Veterinary Parasitology 267, 32–41.

Mazeri, S., Rydevik, G., Handel, I., Bronsvoort, B.M.d., Sargison, N., 2017. Estimation of the impact of Fasciola hepatica infection on time taken for UK beef cattle to reach slaughter weight. Sci. Rep. 7, 7319.

Mehmood, K., Zhang, H., Sabir, A.J., Abbas, R.Z., Ijaz, M., Durrani, A.Z., Saleem, M.H., Ur Rehman, M., Iqbal, M.K., Wang, Y., Ahmad, H.I., Abbas, T., Hussain, R., Ghori, M.T., Ali, S., Khan, A.U., Li, J., 2017. A review on epidemiology, global prevalence and economical losses of fasciolosis in ruminants. Microbial Pathogenesis 109, 253–262.

Mezo, M., González-Warleta, M., Castro-Hermida, J.A., Muiño, L., Ubeira, F.M., 2011. Association between anti-F. hepatica antibody levels in milk and production losses in dairy cows. Veterinary Parasitology 180, 237–242.

Molina-Hernández, V., Mulcahy, G., Pérez, J., Martínez-Moreno, Á., Donnelly, S., O’Neill, S.M., Dalton, J.P., Cwiklinski, K., 2015. Fasciola hepatica vaccine: we may not be there yet but we’re on the right road. Veterinary parasitology 208, 101–111.

Molina, E.C., 2005. Comparison of host-parasite relationships of Fasciola gigantica infection in cattle (Bos taurus) and swamp buffaloes (Bubalis bubalis)’. James Cook University,

Molina, E.C., Gonzaga, E.A., Lumbao, L.A., 2005. Prevalence of infection with Fasciola gigantica and its relationship to carcase and liver weights, and fluke and egg counts in slaughter cattle and buffaloes in Southern Mindanao, Philippines. Tropical Animal Health and Production 37, 215–221.

Novobilský, A., Höglund, J., 2015. First report of closantel treatment failure against Fasciola hepatica in cattle. International Journal for Parasitology: Drugs and Drug Resistance 5, 172–177.

Nyirenda, S.S., Sakala, M., Moonde, L., Kayesa, E., Fandamu, P., Banda, F., Sinkala, Y., 2019. Prevalence of bovine fascioliasis and economic impact associated with liver condemnation in abattoirs in Mongu district of Zambia. BMC veterinary research 15, 33–33.

Paczkowski, M.J., 2005. Effects of experimental fascioliasis on puberty and comparison of mounting activity by radiotelemetry in pubertal and gestating beef heifers. Texas A&M University,

Randall, W.F., Bradley, R.E., 1980. Effects of hexachlorethane on the milk yields of dairy cows in north Florida infected with Fasciola hepatica. American Journal of Veterinary Research 41, 262–263.

Reid, J.F., Doyle, J.J., Armour, J., Jennings, F.W., 1972. Fasciola hepatica infection in cattle. Veterinary Record 90, 486–487.

Roseby, F.B., 1970. The effect of fasciolosis on the wool production of Merino sheep. Australian Veterinary Journal 46, 361–365.

Ross, J.G., Dow, C., 1966. The problem of acute fascioliasis in cattle. Veterinary Record 78, 670.

Sanchez-Vazquez, M.J., Lewis, F.I., 2013. Investigating the impact of fasciolosis on cattle carcase performance. Veterinary Parasitology 193, 307–311.

Sanchez, J., Dohoo, I., Carrier, J., DesCôteaux, L., 2004. A meta-analysis of the milk-production response after anthelmintic treatment in naturally infected adult dairy cows. Preventive Veterinary Medicine 63, 237–256.

Schweizer, G., Braun, U., Deplazes, P., Torgerson, P.R., 2005. Estimating the financial losses due to bovine fasciolosis in Switzerland. Veterinary Record 157, 188–193.

Skuce, P.J., Zadoks, R.N., 2013. Liver fluke – a growing threat to UK livestock production. Cattle Practice 21, 138–149.

Stewart, G.B., Côté, I.M., Rothstein, H.R., Curtis, P.S., 2013. First steps in beginning a meta-analysis, In: Koricheva, J., Gurevitch, J., Mengersen, K. (Eds.) Handbook of Meta-Analysis in Ecology and Evolution. Princeton University Press, Princeton, pp. 27–36.

Sykes, A.R., Coop, R.L., Rushton, B., 1980. Chronic subclinical fascioliasis in sheep: effects on food intake, food utilisation and blood constituents. Research in Veterinary Science 28, 63–70.

Toet, H., Piedrafita, D.M., Spithill, T.W., 2014. Liver fluke vaccines in ruminants: strategies, progress and future opportunities. International Journal for Parasitology 44, 915–927.

Twomey, A.J., Carroll, R.I., Doherty, M.L., Byrne, N., Graham, D.A., Sayers, R.G., Blom, A., Berry, D.P., 2018. Genetic correlations between endo-parasite phenotypes and economically important traits in dairy and beef cattle. J Anim Sci 96, 407–421.

Wamae, L.W., Hammond, J.A., Harrison, L.J.S., Onyango-Abuje, J.A., 1998. Comparison of production losses caused by chronic Fasciola gigantica infection in yearling Friesian and Boran cattle. Tropical Animal Health and Production 30, 23–30.

Wiedosari, E., Hayakawa, H., Copeman, B., 2006. Host differences in response to trickle infection with Fasciola gigantica in buffalo, Ongole and Bali calves. Tropical Animal Health and Production 38, 43–53.

