## Supporting Information for "The influence of liver fluke infection on production in sheep and cattle: a meta-analysis"

Adam D. Hayward, Philip J. Skuce and Tom N. McNeilly

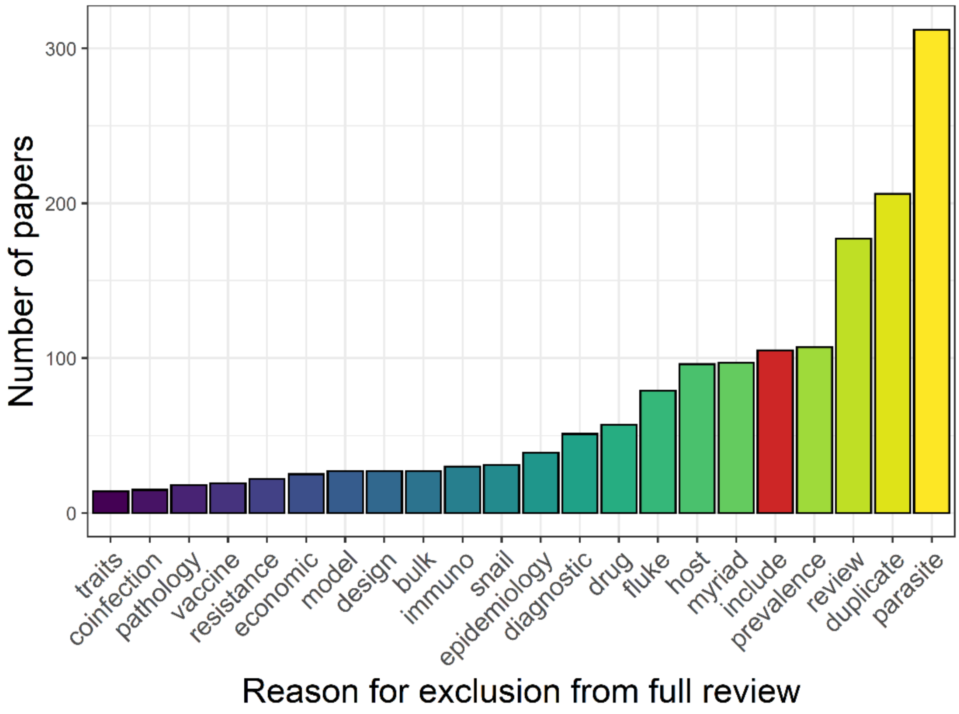


**Figure S1:** Summary of reasons for exclusion from full review of the 1582 papers that resulted from our literature search.

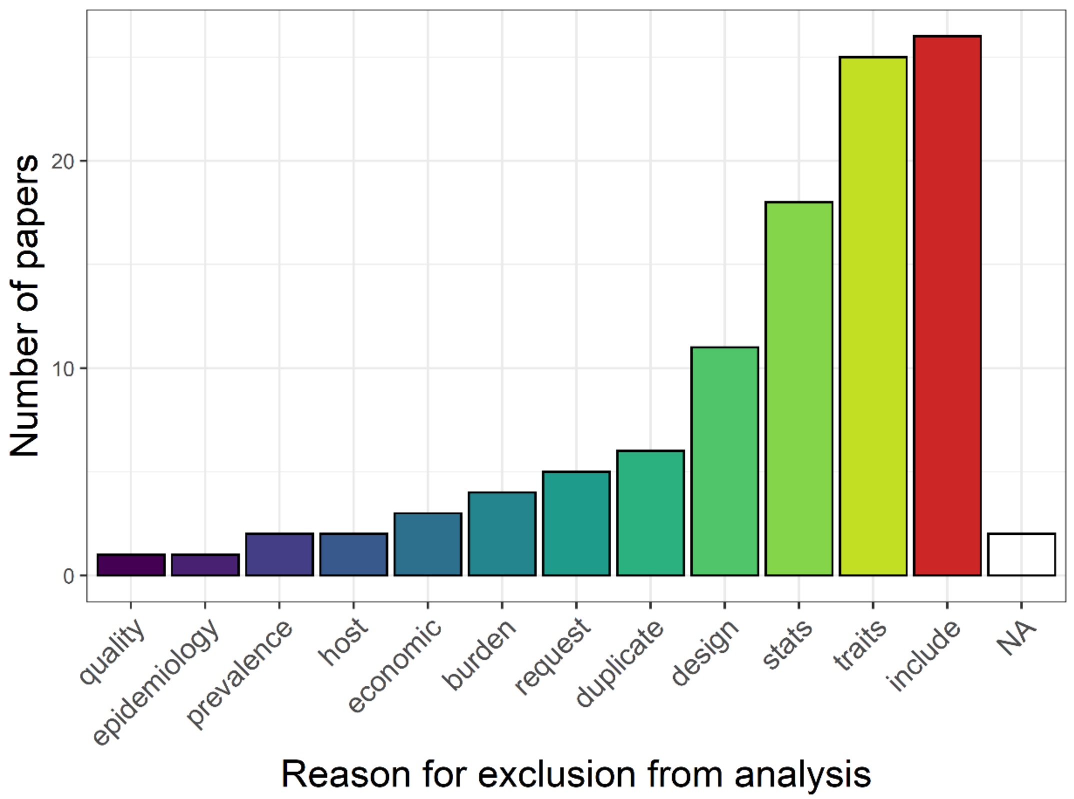


**Figure S2:** Summary of reasons for exclusion from the meta-analysis of the 106 papers that passed our initial screening. “NA” are papers that we were unable to source.

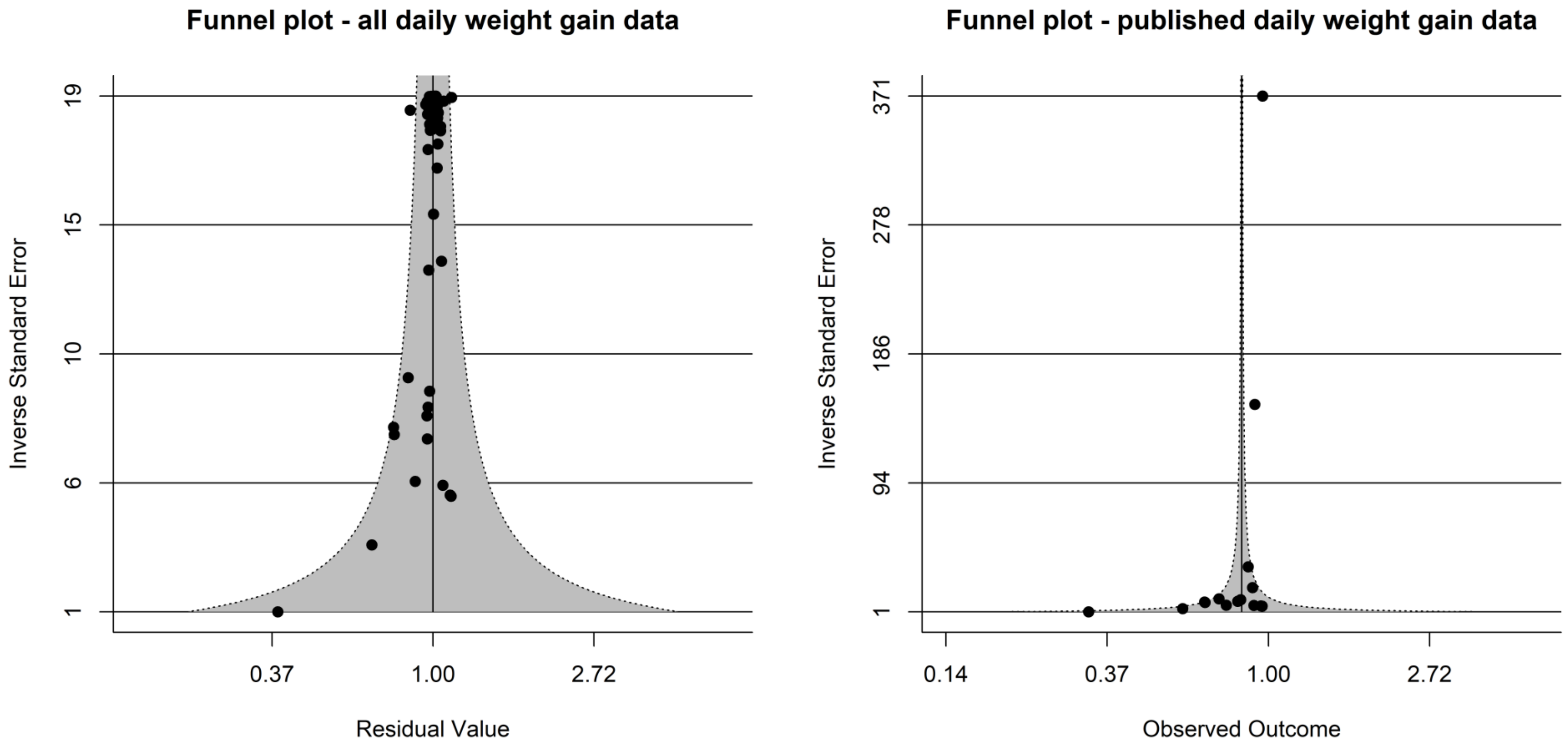


**Figure S3.** Funnel plots showing meta-analytic residuals plotted against inverse standard error of effect sizes for daily weight gain, for both all daily weight gain data (N = 77) and published data (N = 16).

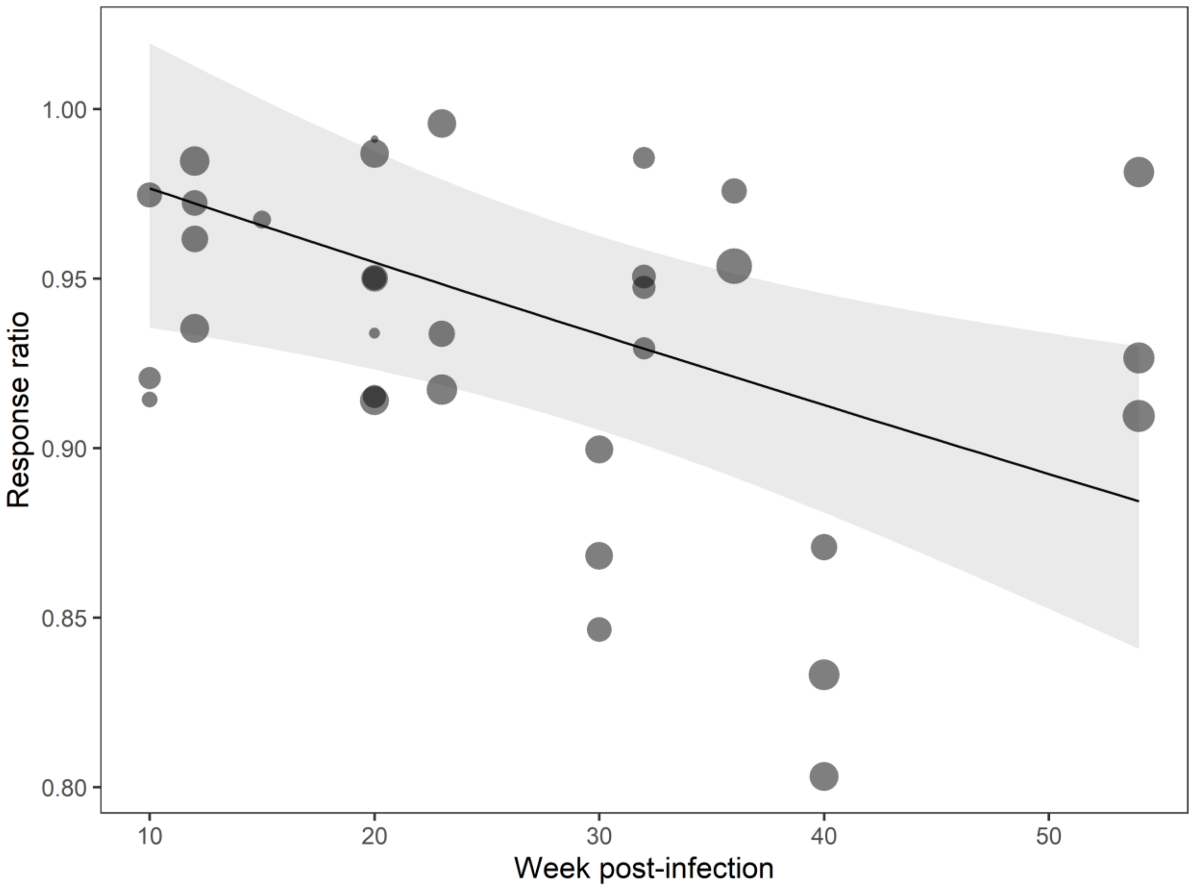


**Figure S4:** The estimated effect of week post-infection on the effect of fluke infection on live weight. Points show raw response ration values, with points sized so that larger points have lower variance; line and shaded area show estimates from a meta-regression with the moderator of week post-infection. A negative slope suggests that infected animals perform less well compared to uninfected animals as the study progresses.

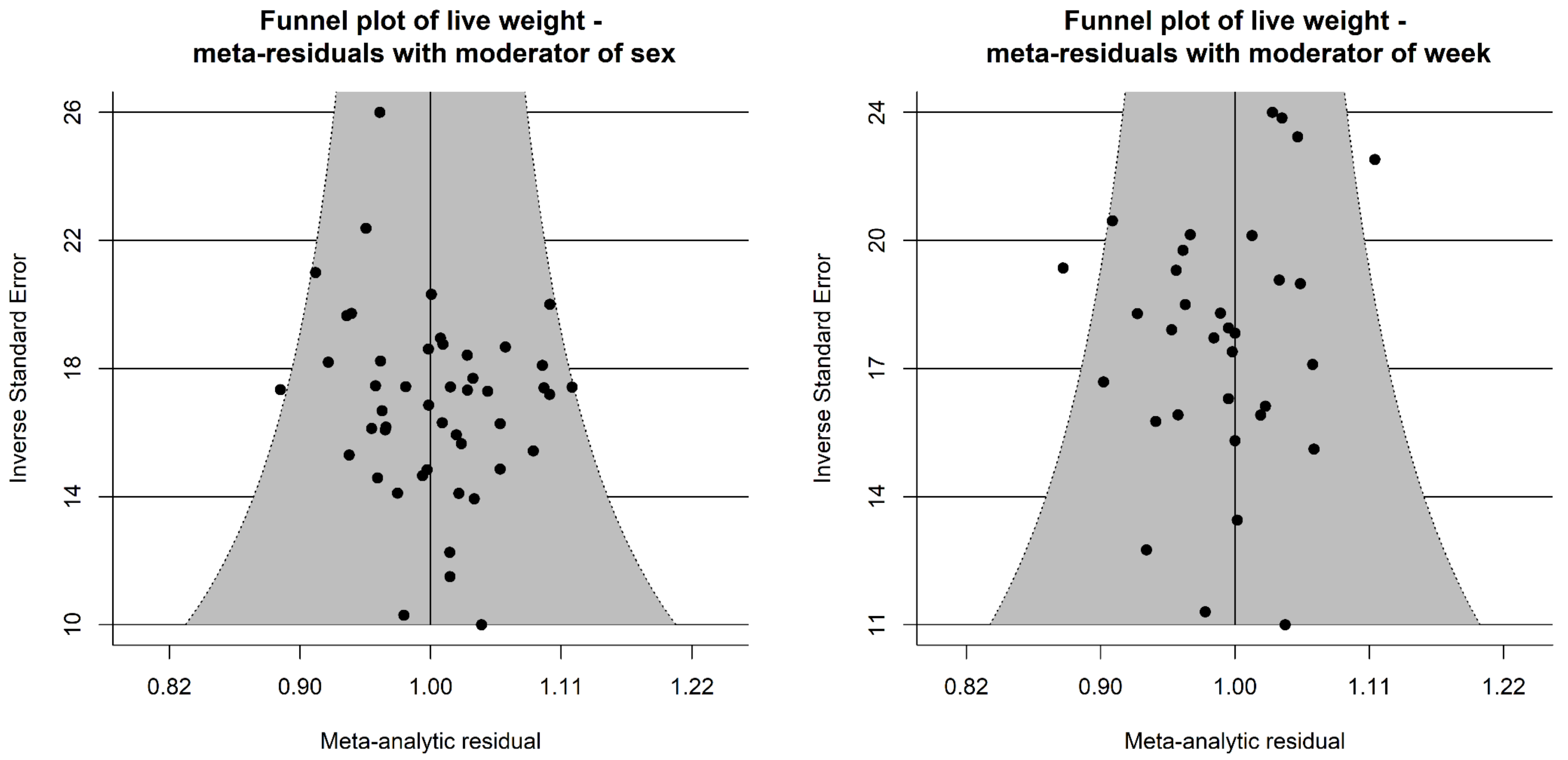


**Figure S5.** Funnel plots showing meta-analytic residuals plotted against inverse standard error of effect sizes for live weight, for both the complete data set including the moderator of sex (N = 47) and the reduced data set with the moderator of week (N = 33).

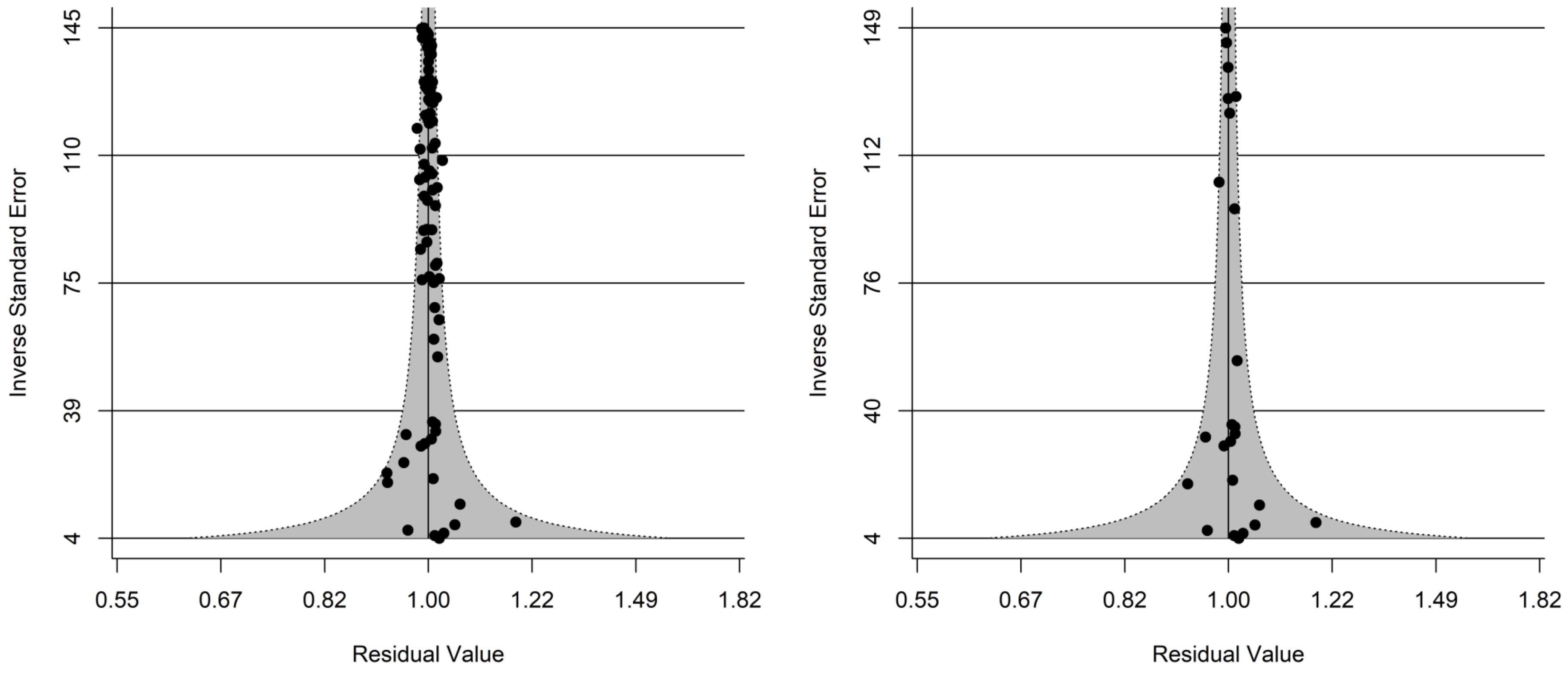


**Figure S6.** Funnel plots showing meta-analytic residuals plotted against inverse standard error of effect sizes for carcass weight, for both the complete data set (N = 84) and the published data (N = 24).

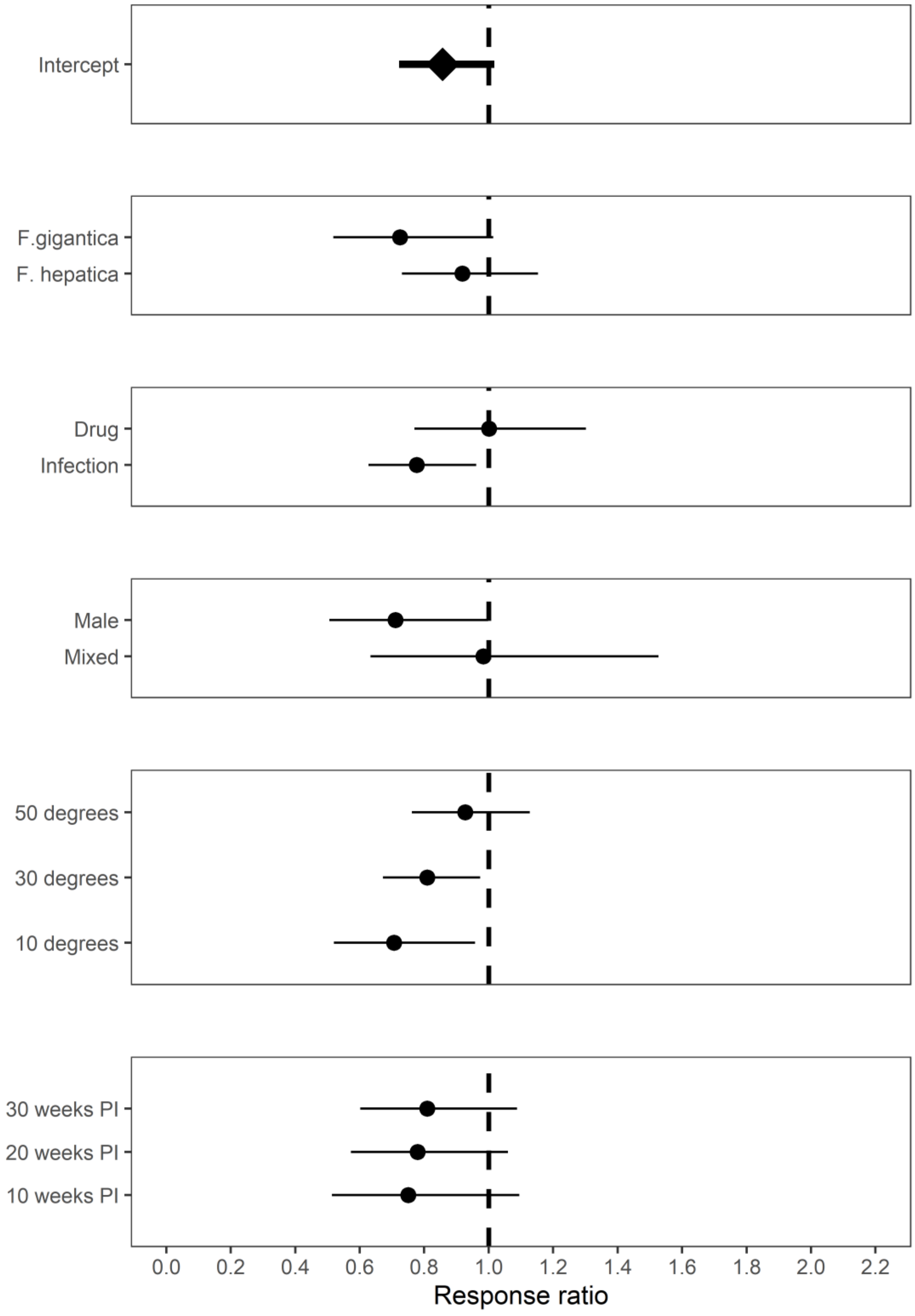


**Figure S7:** Estimated response ratio (RR) from meta-regression analysis of fluke-infected and uninfected animals for total weight gain. The plot shows the estimated mean effect of the global effect on infection, plus the effect of infection in different moderators, with 95%CI. A response ratio of 1 indicates equal performance in infected and uninfected animals; values <1 indicate poorer performance in infected animals.

**Table S1:** Results from meta-analysis of total weight gain. 'N' is the number of effect sizes for each trait. 'Global' indicates the overall effect size from the initial random effects model, while the subsequent moderator effects are from mixed-effects models where each moderator was fitted in turn. 'Effect sizes' and 'Studies' varies because information was not available for some of the effect sizes. '$Q_M$' and the associated 'DF' and 'P' values refer to the Wald-type test for significant differences in effect size due to moderator, while 'z' and 'P' values refer to tests for differences from zero for different levels of each moderator.

| Moderator | N | Studies | $Q_M$ | DF | P | Level | Estimate (logRR) | Lower 95%CI | Upper 95%CI | z | P |
|---|---|---|---|---|---|---|---|---|---|---|---|
| Global | | | | | | | -0.1541 | -0.3258 | 0.0176 | -1.76 | 0.079 |
| Parasite | 18 | 8 | 1.30 | 1 | 0.254 | *F. gigantica* | -0.3218 | -0.6579 | 0.0143 | -1.88 | 0.061 |
| | | | | | | *F.hepatica* | -0.0854 | -0.3132 | 0.1423 | -0.74 | 0.462 |
| Experiment | 18 | 8 | 2.17 | 1 | 0.141 | Drug | 0.0011 | -0.2614 | 0.2637 | 0.01 | 0.993 |
| | | | | | | Infection | -0.2528 | -0.4658 | -0.0397 | -2.33 | 0.020 |
| Sex | 14 | 6 | 1.32 | 1 | 0.251 | Male | -0.3416 | -0.6817 | -0.0016 | -1.97 | 0.049 |
| | | | | | | Mixed | -0.0164 | -0.4561 | 0.4234 | -0.07 | 0.942 |
| Latitude | 18 | 8 | 2.25 | 1 | 0.133 | Continuous | 0.0068 | -0.0021 | 0.0157 | 1.50 | 0.133 |
| Week | 14 | 6 | 0.27 | 1 | 0.605 | Continuous | 0.0038 | -0.0105 | 0.0180 | 0.52 | 0.605 |

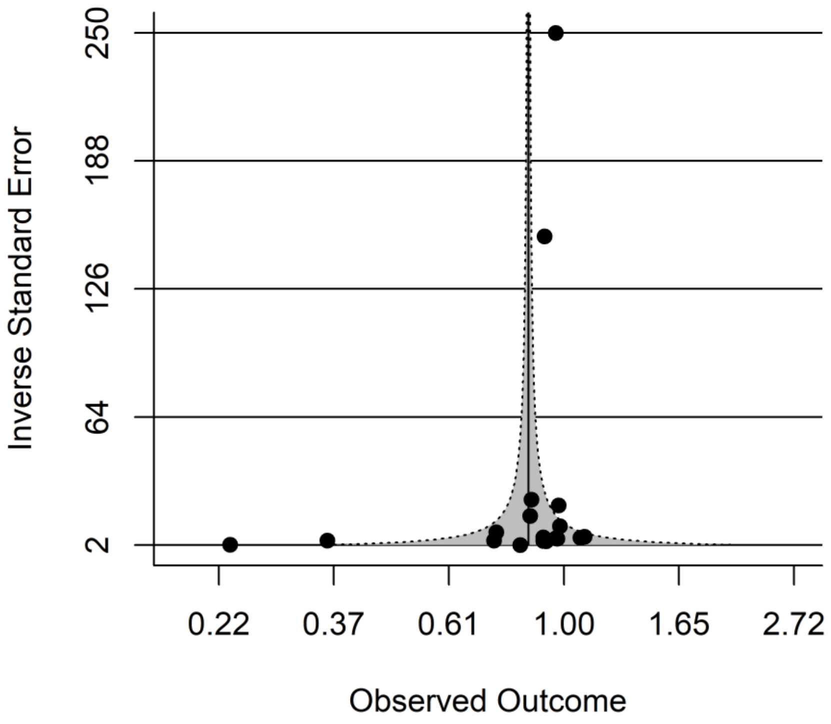


**Figure S8.** Funnel plot showing meta-analytic effect size plotted against inverse standard error of effect sizes for total weight gain.

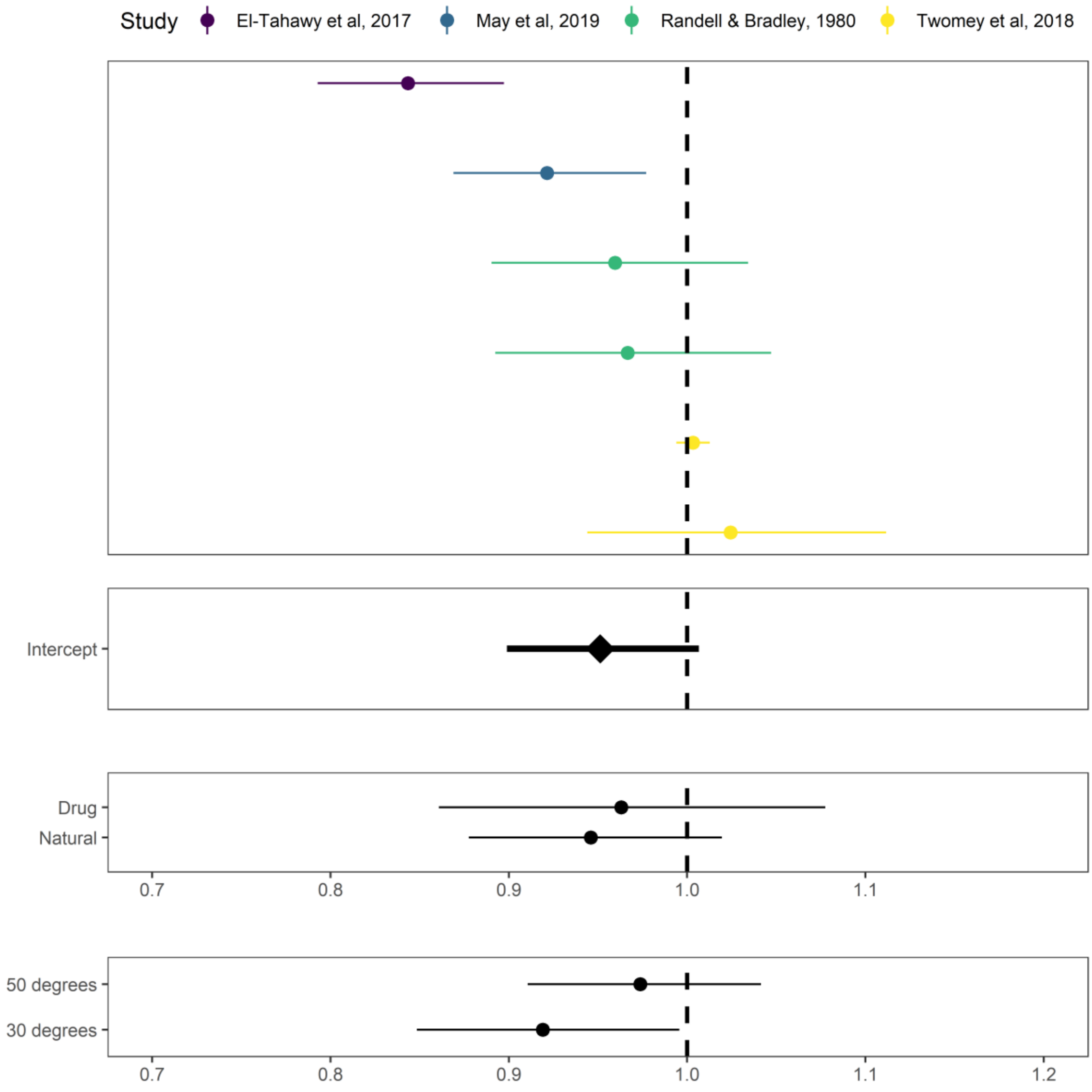


**Figure S9:** Estimated response ratio (RR) from meta-regression analysis of fluke-infected and uninfected animals for milk production. The forest plot (top) shows each of the six effect size estimates that our analyses were based on. The proceedings plots show estimated mean effect of the global effect on infection, plus the effect of infection in different moderators, with 95%CI. A response ratio of 1 indicates equal performance in infected and uninfected animals; values <1 indicate poorer performance in infected animals.

| Moderator | N | Studies | $Q_M$ | DF | P | Level | Estimate (logRR) | Lower 95%CI | Upper 95%CI | z | P |
|---|---|---|---|---|---|---|---|---|---|---|---|
| Global | 6 | 4 | | | | | -0.0500 | -0.1065 | 0.0066 | -1.73 | 0.083 |
| Experiment | 6 | 4 | 0.07 | 1 | 0.794 | Drug | -0.0376 | -0.1499 | 0.0746 | -0.66 | 0.511 |
| | | | | | | Natural | -0.0556 | -0.1305 | 0.0193 | -1.45 | 0.146 |
| Latitude | 6 | 4 | 1.32 | 1 | 0.251 | Continuous | 0.0029 | -0.0020 | 0.0078 | 1.15 | 0.251 |

**Table S2:** Results from meta-analysis of milk production. ‘N’ indicates the number of effect sizes available for each modifier. ‘Global’ indicates the overall effect size from the initial random effects model, while the subsequent moderator effects are from mixed-effects models where each moderator was fitted in turn. ‘Effect sizes’ and ‘Studies’ varies because information was not available for some of the effect sizes. ‘$Q_M$’ and the associated ‘DF’ and ‘P’ values refer to the Wald-type test for significant differences in effect size due to moderator, while ‘z’ and ‘P’ values refer to tests for differences from zero for different levels of each moderator.

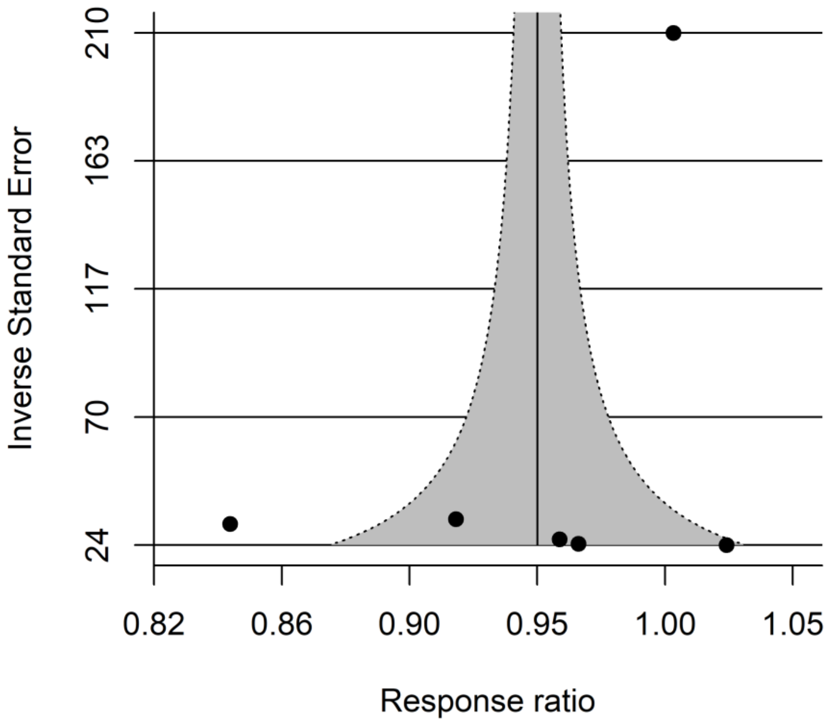


**Figure S10.** Funnel plot showing meta-analytic effect size plotted against inverse standard error of effect sizes for milk production.
